# A myosin chaperone, UNC-45A, is a novel regulator of intestinal epithelial barrier integrity and repair

**DOI:** 10.1101/2021.11.10.467974

**Authors:** Susana Lechuga, Alexander X. Cartagena-Rivera, Afshin Khan, Bert I. Crawford, Vani Narayanan, Daniel E. Conway, Jaakko Lehtimäki, Pekka Lappalainen, Florian Rieder, Michelle S. Longworth, Andrei I. Ivanov

**Affiliations:** Department of Inflammation and Immunity, Lerner Research Institute, Cleveland Clinic Foundation, Cleveland, OH; Section on Mechanobiology, National Institute of Biomedical Imaging and Bioengineering, National Institutes of Health, Bethesda, MD; Department of Biomedical Engineering, Virginia Commonwealth University, Richmond, VA; Institute of Biotechnology and Helsinki Institute of Life Sciences, University of Helsinki, Helsinki, Finland; Department of Gastroenterology, Hepatology and Nutrition, Digestive Diseases and Surgery Institute, Cleveland Clinic Foundation, Cleveland, OH

## Abstract

The actomyosin cytoskeleton serves as a key regulator of the integrity and remodeling of epithelial barriers by controlling assembly and functions of intercellular junctions and cell-matrix adhesions. While biochemical mechanisms that regulate activity of non-muscle myosin II (NM-II) in epithelial cells have been extensively investigated, little is known about assembly of the contractile myosin structures at the epithelial adhesion sites. UNC-45A is a cytoskeletal chaperone that is essential for proper folding of NM II heavy chains and myofilament assembly. We found abundant expression of UNC-45A in human intestinal epithelial cell (IEC) lines and in the epithelial layer of the normal human colon. Interestingly, protein levels of UNC-45A were decreased in colonic epithelium of patients with ulcerative colitis. CRISPR/Cas9-mediated knock-out of UNC-45A in HT-29 and SK-CO15 IEC disrupted epithelial barrier integrity, impaired assembly of epithelial adherence and tight junctions and attenuated cell migration. Consistently, decreased UNC-45 expression increased permeability of the *Drosophila* gut *in vivo*. The mechanisms underlying barrier disruptive and anti-migratory effects of UNC-45A depletion involved disorganization of the actomyosin bundles at epithelial junctions and the migrating cell edge. Loss of UNC-45A also decreased contractile forces at epithelial junctions and matrix adhesions. Expression of deletion mutants revealed roles for the myosin binding domain of UNC-45A in controlling IEC junctions and motility. Our findings uncover a novel mechanism that regulates integrity and restitution of the intestinal epithelial barrier, which may be impaired during mucosal inflammation.

## Introduction

Formation of intercellular junctions represents a key molecular event that is required for epithelial cell differentiation, apico-basal polarization and establishment of the paracellular barrier ^1^. It is also essential for tissue morphogenesis by enabling the orchestrated movement, expansion and folding of epithelial sheets ^2,3^. While cell-cell engagement results in the assembly of several adhesive structures, the apically located tight junctions (TJ) and adherens junctions (AJ) are arguably the most important regulators of epithelial barriers and homeostasis ^4–7^. TJ and AJ are elaborate multiprotein complexes that include transmembrane adhesive proteins connected to various intracellular scaffolds. Transmembrane AJ (e.g., E-cadherin, nectins) and TJ (claudins, occludin, junctional adhesion molecule A, etc.) proteins at the opposing plasma membranes mediate cell-cell adhesions and epithelial barrier formation ^4–7^. The intracellular scaffolds that include members of a ‘*zonula occludens*’ (ZO) protein family at TJ and different catenin proteins at AJ, orchestrate assembly, clustering and remodeling of transmembrane adhesive modules of epithelial junctions ^4–7^.

AJ and TJ functions in regulating epithelial barriers and morphogenesis depend on their ability to sense and transduce mechanical forces ^8,9^. Such ability is determined by coupling of junctional complexes to the cortical actomyosin cytoskeleton. In simple differentiated epithelia, the most noticeable cytoskeletal structure is a circumferential F-actin belt that is directly associated with AJ and TJ ^10,11^. This belt is enriched with a key actin motor protein, non-muscle myosin II (NM-II). It has been firmly established that NM-II activity mediates both passive tension, required for maintenance of epithelial barrier integrity and contractile forces that mediate junctional remodeling (disassembly and reassembly) ^8,9,12^.

Superresolution microscopy imaging revealed highly ordered periodic distribution of NM-II clusters in the perijunctional F-actin belt of epithelial tissues that resembles the sarcomeric organization of actomyosin units in muscle cells ^10,13^. While such high-order organization is likely to be important for the spatio-temporal NM-II interactions with actin filaments and force generation, little is known about the mechanisms regulating the ordered, myofibril-like assembly of contractile NM-II units in the perijunctional actomyosin belt. Incorporation of myosin II molecules into myofibrils is known to be orchestrated with the folding of the myosin motor domain and requires cytosolic molecular chaperones ^14^. UNC-45 belongs to the evolutionary conserved UCS (**U**NC-45/**C**ro1/**S**he4p) protein family, and is the most studied myosin chaperone regulating a variety of actomyosin driven processes in high eukaryotes ^14,15^. Vertebrate organisms express two different paralogs, UNC-45A and UNC-45B, which have ~55% of the sequence identity. Whereas UNC-45A is ubiquitously expressed in different tissues, UNC-45B is mainly expressed in muscle cells ^16^. UNC-45 contains three different domains: the N-terminal tetratricopeptide (TPR) domain, the central region and the C-terminal USC domain ^14,15,17^. The USC domain directly binds to myosin II motor and regulates myosin folding ^18–21^. The TPR domain interacts with Hsp90 and Hsp70, which serve as co-chaperones assisting UNC-45-mediated myosin folding ^19,20,22^. Recent studies in *C. elegans* revealed that intermolecular interactions between the TPR and USC domains are essential for formation of UNC-45 oligomers ^20,21^. There, oligomerization of this chaperone into linear molecular arrays promotes the assembly of folded myosin II molecules into highly ordered myofilaments. Notably, UNC-45A has been recently shown to bind and destabilize microtubules ^23–25^. Such microtubule-destabilizing activity requires the TRP domain but appears to be independent of myosin binding and folding by UNC-45A ^23^.

The majority of functional studies of UNC-45 were conducted in model organisms such as *C. elegans* and *Drosophila* and focused on muscle development and contractility ^14,15,17^. However, little is known about the functions of this chaperone in mammalian non-muscle cells. Previously described phenotypes of UNC-45A depletion in these systems include attenuation of growth cone development in neurons ^26^, as well as modulation of proliferation and motility of different cancer cells ^27–30^. However, UNC-45A involvement in the regulation of epithelial barrier integrity and remodeling has remained largely unknown.

We are filling these crucial knowledge gaps by reporting that UNC-45A acts as a positive regulator of barrier properties both in the human intestinal epithelial monolayers as well as in the *Drosophila* gut. Such barrier enhancing activity of this chaperone involves controlling the assembly of apical junctions and the perijunctional actomyosin cytoskeleton. We also observed that UNC-45A regulates the repair of injured intestinal epithelium by modulating cell-matrix adhesions and promoting wound healing. Together, our findings signify this myosin chaperone as a novel regulator of the establishment and remodeling of epithelial barriers.

## Materials and Methods

### Antibodies and other reagents

The following monoclonal (mAb) and polyclonal (pAb) antibodies were used to detect apical junction, cytoskeletal and matrix adhesion proteins: E-cadherin (610181), β-catenin (610154), mAbs (BD Biosciences, San Jose, CA); anti-cleaved PARP (9541), cleaved caspase3 (9664), NM-IIC (3405) pAbs (Cell Signaling, Beverly, MA); anti-ZO-1 (402200), phosphorylated paxillin (44722G) pAbs (Thermo Fisher Scientific, Waltham, MA); anti-occludin (13409), α-catenin (12831), GFP (50430) and anti-UNC-45A (19564) pAbs (Proteintech, Rosemont, IL); anti-E-cadherin (AF748) pAb (R & D System, Minneapolis, MN); NM IIA (909801) pAb (Biolegend, San Diego, CA); α-tubulin (T6199) mAb (Millipore-Sigma, San Louis, MO), anti-NM-IIA (H00004627-M03) mAb (Abnova, Walnut, CA). A conformation sensitive α18 α-catenin rat mAb was provided by Dr. Akira Nagafuchi, (Nara Medical University, Japan) Alexa Fluor-488-conjugated donkey anti-rabbit, Alexa Fluor-555-conjugated donkey anti-mouse secondary antibodies were obtained from Thermo-Fisher. Horseradish peroxidase-conjugated goat anti-rabbit and anti-mouse secondary antibodies were obtained from Bio-Rad Laboratories. Antibodies used for immunoblotting of *Drosophila* lysates included `β`-tubulin (Cell Signaling, 2146) and Unc-45 (gift from Dr. Sanford Bernstein, San Diego State University ^31^. Others reagents were obtained from Millipore-Sigma.

### Drosophila Stocks

Standard laboratory strains *w*^*1118*^ and *unc-45*^*03692/+*^ were obtained from Bloomington *Drosophila* Stock Center. *unc-45*^*EY03043–33/+*^ was a generous gift from the Bernstein laboratory at San Diego State University.

### Cell Culture and calcium switch

SK-CO15 human colonic epithelial cells ^32^ were a gift from Dr. Enrique Rodriguez-Boulan (Weill Medical College of Cornell University, New York, NY). HT-29cf8, a well-differentiated clone of HT-29 cells ^33,34^ was provided by Dr. Judith M. Ball (College of Veterinary and Biomedical Sciences, Texas A&M University, College Station, TX). Caco-2, T84, Colo320 and SW620 cell lines were obtained from the American Type Culture Collection (Manassas, VA). HT-29cf8 and SK-CO15 cell lines were authenticated by the small tandem repeat profiling by the LabCorp (Burlington, NC). All cultured cells were mycoplasma free according to the mycoplasma PCR detection assay (PromoCell, Heidelberg, Germany). HT-29cf8, and SK-CO15 cells were cultured in DMEM medium supplemented with 10% fetal bovine serum (FBS), HEPES, non-essential amino acids, and penicillin-streptomycin antibiotic. The cells were seeded on collagen-coated Transwell filters (Corning Incorporated, Corning, NY), or glass coverslips for permeability measurements and immunolabeling experiments and on 6-well plastic plates for other functional and biochemical studies. The calcium switch assay was performed as previously described ^35–37^. Briefly, well-differentiated SK-CO15 cell monolayers were subjected to overnight extracellular calcium depletion by incubating in the Eagle’s minimum essential medium for suspension culture (Thermo Fisher Scientific) supplemented with 5 μM CaCl_2_, 10 mM HEPES and 10% dialyzed FBS, pH 7.4. To induce junctional reassembly, the cells were returned to a normal cell culture medium with high (~ 1.8 mM) calcium for the indicated time.

### Primary human intestinal epithelial cells

Deidentified full thickness intestinal surgical resection specimen obtained from patients with active Crohn’s Disease, ulcerative colitis and non-IBD controls (diverticulosis or non-affected margins of resected colorectal cancer) were provided by the Biorepository Core of the Cleveland Digestive Diseases Research Core Center. Primary colonic epithelial cells were isolated based on a previously described protocol ^38^. Briefly, resected samples were cut into approximately 1.5 cm long strips. Colonic epithelial cells were isolated by incubating the strips in HEPES-buffered Hanks’ balanced salt solution (HBSS) containing 1 mM EDTA (pH 7.4) under constant agitation for 3-h at room temperature. Detached epithelial cells were centrifuged for 10 min at 3000 rpm. Cells were washed once in ice-cold HBSS, resuspended into RIPA buffer and homogenized.

### CRISPR-Cas9 mediated knockout of UNC-45A and generation of UNC-45A overexpressing IEC

A stable knockout of UNC-45A in HT-28cf8 and SK-CO15 IEC was carried out using a CRISPR-Cas9 technology. The guide oligonucleotide sequences used for knocking out UNC-45A in HT-29cf8 cells were: sgRNA1, forward 5’-CACCGTCTCTAGGGCTTGGCTCCGC-3’ and reverse 5’-AAACGCGGAGCCAAGCCCTAGAGAC-3’; and sgRNA2 forward 5’-CACCGGGGGCGTCGCGTCCAGACCC-3’ and reverse 5’-AAACGGGTCTGGACGCGACGCCCCC-3’. For knocking out UNC-45A in SK-CO15 cells the sgRNA forward 5’-CACCGGCAGGCTTCTCAGAACCTGG-3’ and reverse 5’-AAACCCAGGTTCTGAGAAGCCTGCC-3’ were used. The guide oligonucleotides were phosphorylated, annealed and cloned into the BsmBI site of a lentiCRISPR v2 vector (Addgene, 52961) according to a published protocol ^39,40^. Obtained constructs were verified by sequencing. Transfer plasmids possessing annealed guide oligonucleotides were transformed into recombination-deficient Stbl3 bacteria and amplified plasmids were isolated from the bacteria using Qiagen Midi prep plasmid isolation kit. Lentiviruses were produced by transfecting HEK-293T cells with the transfer lentiCRISPR v2 plasmids and packaging plasmids pLTR-G (Addgene, 17532) and pCD/NL-BH*DDD (Addgene, 17531). Viral supernatants were collected 48 h and 72 h after transfection and used to infect IEC. After 24 h of the transduction, the lentivirus-containing medium were replaced with fresh cell culture medium containing puromycin (10 μg/ml) and puromycin-resistant cells were collected after 7 day selection. After puromycin selection, single clone isolation was performed in SK-CO15 cells by using single cell dilution protocol and visually screening 98-wells plates for single cell identification. When colonies reach 70% confluence, cells were harvested and the UNC45A depletion was analyzed by immunoblotting.

SK-CO15 cell lines were generated to stable express either full length UNC-45A (UNC-45A-GFP), its N-terminal truncated ΔTPR-GFP (residues 147-971), or C-terminally truncated ΔUCS-GFP (residues 1-539). Cloning of the full length and deletion mutant UNC-45A constructs into pEGFP-N3 vector (Clontech Laboratories) has been previously described ^30^. IEC were transfected using Lipofectamine 2000 (Thermo Fisher Scientific). 48 h after transfection, fresh medium containing G418 antibiotic (1mg/ml) was added. After 7 days of antibiotic selection, GFP expressing cells were sorting using FACSAria™ fusion flow cytometer (BD Bioscience). Briefly, cultured cells were harvested and resuspended in PBS with 1% of FBS. After sorting cells were collected in DMEM with 10% FBS and the overexpression of full length and deletion mutant UNC-45A was analyzed by immunoblotting.

### Measurement of epithelial barrier permeability

Transepithelial electrical resistance (TEER) of HT-29cf8, and SK-CO15 intestinal epithelial cell monolayers cultured on collagen-coated transwell filters (Corning Incorporated) was measured using an EVOM2 Volt Ohm Meter (World Precision Instruments, Sarasota, FL). The resistance of cell-free collagen-coated filters was subtracted from each experimental point. A transmonolayer dextran flux assay was performed as previously described ^41,42^. IEC monolayers cultured on transwell filters were apically exposed to 1 mg/ml of FITC-labeled dextran (4,000 Da) in HEPES-buffered Hanks’ balanced salt solution (HBSS). After 120 min of incubation, samples were collected from the lower chamber and FITC fluorescence intensity was measured using a SpectraMax M2 plate reader (Molecular Devices, San Jose, CA), at excitation and emission wavelengths 485 nm and 544 nm, respectively. The amount of FITC-dextran translocated across the epithelial cell monolayer was calculated based on a calibration curve using Prism 9.10 software (GraphPad, La Jolla, CA).

### Immunoblotting analysis and immunoprecipitation

IEC monolayers were scraped and homogenized using a Dounce homogenizer in RIPA buffer (20 mM Tris, 50 mM NaCl, 2 mM EDTA, 2 mM EGTA, 1% sodium deoxycholate, 1% Triton X-100 (TX-100), and 0.1% SDS, pH 7.4) containing protease inhibitor cocktail and phosphatase inhibitor cocktails 2 and 3 (Millipore-Sigma). 15 adult *Drosophila* males and females of different genotypes were harvested at 10 days post-enclosure, crushed in high salt lysis buffer (300 mM NaCl, 50 mM Tris pH 7.5, 1 mM EDTA, 0.1%Triton, 10% glycerol, 1 mM DTT, Complete Mini EDTA-free protease tablets (Roche), and incubated on ice for 15 minutes. The obtained total cell lysates were cleared by centrifugation (20 min at 14,000 x g), diluted with 2x SDS sample loading buffer, and boiled. SDS-polyacrylamide gel electrophoresis was conducted using a standard protocol with equal amounts of total protein (10 μg or 20 μg of protein from human cells or 30 μg of protein per genotype for *Drosophila* extracts) loaded per each lane. The separated proteins were transferred to nitrocellulose membranes and the membranes were blocked with 5% non-fat milk. The blocked membranes were incubated overnight with primary antibodies, and then exposed to HRP-conjugated secondary antibodies for 1 h. The labeled proteins were visualized using a standard enhanced chemiluminescence solution and X-ray films. For immunoblotting of *Drosophila* proteins, bands were quantified from films, using a BioRad ChemiDocTM XRS+ Molecular Imager and Image LabTM software, with “Protein Gel” settings, using identically sized windows to calculate Absolute volumes, as directed by the manufacturer. All bands were first normalized to respective loading controls, prior to comparison between lanes.

Immunoprecipitation (IP) was performed as previously described ^43^. Briefly, cells were homogenized in the IP buffer (50 mM 1,4-piperrazinediethanesulfonic acid, 50 mM HEPES, 1 mM EDTA, 2 mM MgSO_4_, 1% TX-100, and 0.5% Igepal, pH 7.0) containing protease inhibitor and phosphatase inhibitor cocktails 2 and 3. After centrifuged, supernatants were precleared with protein A-sepharose beads (GenScript, Piscataway, NJ). Precleared supernatants were then incubated overnight at 4°C with 5μg of anti-NM IIA pAb or control rabbit immunoglobulin. Immunocomplexes were recovery by incubation with protein A-sepharose beads for 2h at 4°C. After 3 washes with IP buffer, beads were boiled with 2x SDS sample loading buffer and pelleted by centrifugation. 20 μl of supernatants were loaded into SDS-polyacrylamide gel electrophoresis using a standard protocol as described above.

### Immunofluorescence labeling, confocal microscopy and image analysis

To visualize the structure of epithelial junctions and localization of UNC-45A and NM II heavy chains, cultured IEC monolayers were fixed and permeabilized with 100% methanol for 20 min at −20 °C. To visualize the actin cytoskeleton, cells were fixed with 4% PFA for 20 min and permeabilized with 0.5% Triton X-100 for 5 min at room temperature. Fixed samples were blocked for 60 min in HBSS containing 1% bovine serum albumin, followed by a 60-min incubation with primary antibodies. Then, samples were washed and incubated with Alexa-Fluor-488–conjugated donkey anti-rabbit and Alexa-Fluor-555– conjugated donkey anti-mouse secondary antibodies for 60 min, rinsed with blocking buffer, and mounted on slides with ProLong Antifade mounting reagent (Thermo-Fisher Scientific). For the actin cytoskeleton labeling, fixed/permeabilized cells were incubated for 1 h with Alexa-Fluor-488-conjugated phalloidin. Immunofluorescence labeled cell monolayers were imaged using a Leica SP8 confocal microscope (Wentzler, Germany). The Alexa Fluor 488 and 555 signals were acquired sequentially in frame-interlace mode, to eliminate cross talk between channels.

To quantify the intensity of a dual alpha-catenin and α18-α-catenin immunolabeling, four images were taken per slide and signal intensity of 50 cell-cell contacts in each image were measured by ImageJ software (National Institute of Health, Bethesda, MD) and averaged to produce a single data point. For the quantification of the junctional lengths in the calcium switch experiments, four images were taken per slide and the total length of junctional labeling per image was measured by using ImageJ software. To quantify thickness of the perijunctional myosin belt, four images were taken per slide and the width of 50 cell-cell contacts were measured by ImageJ and averaged to produce a single data point. Quantification of focal adhesions (FA) was performed by counting the number of dots on p-paxillin images using ImageJ. Number of FA in three different images was averaged to produce a single data point.

### Proximity ligation assay

Proximity ligation assay (PLA) was performed by using a DUOLINK kit (Millipore-Sigma) according to the manufacturer’s instructions. Briefly, IEC monolayers were fixed in ice-cold methanol and incubated for 1 h with a DUOLINK blocking solution at 37°C. After two washes, anti-UNC-45A polyclonal and anti-NM-IIA monoclonal antibodies diluted in a DUOLINK antibody diluent were added and incubated for 1 h at 37°C. As a negative control, anti-UNC-45A antibody was added alone, without the anti-NM-IIA antibody. Next, the slides were washed and incubated for 1 h with secondary antibodies (DUOLINK anti-rabbit PLA-plus probe and DUOLINK anti-mouse PLA-minus probe). The labeling was completed by consequent incubation with a ligation solution for 30 min and an amplification solution for 100 min at 37°C. Labeled slides were mounted with the Prolong Antifade mounting medium with DAPI. Images were acquired using a Leica SP8 confocal microscope and processed using ImageJ. For quantification, four images per slide were taken and the total number of dots was measured by using ImageJ software and divided by the number of cells.

### FRET Biosensors

A previously developed and validated E-cadherin FRET-based tension biosensor ^44^ was used to measure E-cadherin mechanical tension. This sensor was expressed in HT-29cf8 cells using adenovirus. The adenovirus was constructed by subcloning the existing E-cadherin tension sensor into the pShuttle-CMV vector (Addgene, 16403), followed by recombination into the pAdEasy1 vector (Addgene, 16400). Adenovirus was prepared using standard protocols ^45^.

HT-29cf8 cells expressing the E-cadherin tension sensor were grown on Transwell inserts (Corning Incorporated). Inserts were cut out, flipped over, and living cells were imaged through glass bottom dishes using a plan-apochromat 40x water immersion NA 1.1 objective lens on an inverted Zeiss LSM 710 laser scanning microscope (Oberkochen, Germany). All the images were acquired at 458 nm excitation wavelength from an Argon laser source. Using the online-unmixing mode, both mTFP1 (donor) and mEYFP (acceptor) channels were collected via spectral unmixing as previously described ^46^. Intensity images were further processed and analyzed using a custom Python code, which involved background subtraction and removal of saturated pixels, as previously described ^46^. The FRET index images were obtained by taking the ratio of the acceptor fluorophore channel (mEYFP) to the donor fluorophore channel (mTFP1), which was then multiplied with the binary image masks that outlined the cell-cell junctions in order to inspect FRET pixels of interest. When comparing multiple groups, all pixels of interest were aggregated following which, an upper and lower bound for the intensity was chosen in order to exclude dim pixels.

### Acoustic frequency modulated atomic force microscopy

Epithelial viscoelastic properties measurements in control and UNC-45A-depleted HT-29cf8 epithelial cell monolayers by noncontact acoustic Frequency Modulation Atomic Force Microscopy (FM-AFM) ^47,48^. HT-29cf8 cells were seeded on glass-bottom petri dishes (Willco Wells) and cultured for 7-10 days to achieve monolayer confluency. FM-AFM experiments were performed utilizing a Bruker Bioscope Catalyst Atomic Force Microscope system (Bruker) mounted on an inverted Axiovert 200M optical microscope (Carl Zeiss) equipped with a Confocal Laser Scanning Microscope 510 Meta (Carl Zeiss), and using a 40x objective lens (0.95 NA, Plan-Apochromat; Carl Zeiss). During experiments, cell monolayers were maintained at 37°C using a heating sample stage (Bruker). A modified AFM microcantilever with an attached 25μm polystyrene bead (Novascan) was used for all FM-AFM measurements. Using the thermal tune fluctuations method built in the AFM system, the calibrated spring constant were between 0.6N/m-0.14N/m. Next, explants were placed on the AFM sample stage and tapping mode was selected. Then, the cantilever tune curve mode was engaged and the cantilever driving frequency was chosen to be the largest frequency peak near the cantilever natural resonance frequency (predetermined by the thermal tune method). After gently engaging the cell monolayer apical surface, the cantilever tune was launched, and the cantilever was positioned at 8μm away from the epithelium. Then, a frequency sweep was recorded, and the cantilever phase lag corrected to be π/2. Next, the acoustically vibrating bead is moved from the 8μm to 1μm gap distance by 500 nm intervals. A frequency sweep is recorded for each interval. Supracellular apical epithelial tension and viscosity calculations were performed using a custom-made MATLAB script (MathWorks). The apical epithelial mechanics model based on lubrication theory for linearized Stokes flow used herein can be accessed for detailed description ^48^.

### Scratch wound assay

IEC were plated into Culture-Insert 3 Well in μ-Dish (Ibidi GmbH, Germany) and allowed to grow to confluence. A 3-well silicone gasket was removed and the bottom of the dish was marked to define the position of the wound. The monolayers were supplied with fresh cell culture medium and images of a cell-free area at the marked region were acquired at 0 h and 24 h after the gasket removal using a Keyence BZ-X710 microscope (Keyence, Itasca, IL). The wound area of three different wounds was measures and the percentage of wound closure was calculated using Image J.

### Boyden chamber migration assay

Boyden chamber migration assay was performed using Transwell® 6.5 mm membrane inserts with 8.0 μm pores (Corning Incorporated). The membrane inserts were coated with 15 μg/cm^2^ of collagen I. Cells were detached from the plate using a TrypLE Express solution (Thermo-Fisher Scientific), resuspended in serum-free medium, and added to the Transwell upper chamber at the density of 5,000 HT-29cf8 cells and 6,000 SK-CO15 cells per chamber. A complete cell culture medium containing 10% FBS as a chemoattractant was added to the lower chamber and cells were allowed to migrate for 16 h at 37°C. Membrane inserts were fixed with methanol and non-migrated cells were removed from the top of the filter using a cotton swab. The cells remained at the bottom of the filter were labeled with DAPI nuclear stain and images using the Keyence BZ-X710 microscope. Three images per membrane insert were taken and number of DAPI-positive cells in each image was counted using the Image J and averaged to yield a single data point.

### Extracellular matrix adhesion assay

Cells were detached from the plate, counted with a T20 automated cell counter (Bio-Rad), and resuspended in the complete medium. Ten thousand (10,000) HT-29cf8 cells and 7,500 SK-CO15 cells were seeded per well in a 24-well plate coated with rat collagen I and were allowed to adhere for 30 min at 37 °C. After incubation, unattached cells were aspirated and the wells were gently washed with HBSS buffer. The attached cells were fixed with methanol and stained using a DIFF stain kit (Electron Microscopy Sciences, Hatfield, PA). Images of adherent cells were captured using the Keyence BZ-X710 microscope. Three images were taken for each well and number of adhered cells in each image was counted using the Image J and averaged to yield a single data point.

### Cell proliferation assay

Cell proliferation was examined by the MTT assay involving the conversion of the water-soluble MTT (3-(4,5-dimethylthiazol-2-yl)-2,5-diphenyltetrazolium bromide) into insoluble formazan. Cells were cultured on 98 well plates, and at different times post-plating, their medium was replaced with 100 μL of fresh culture medium containing 50 μg/ml of MTT. MTT added to 100 μL of medium alone was included as a negative control. Cells were incubated at 37°C for 4 h and the generated formazan was dissolved by adding 50 μL of DMSO with 10 min incubation at 37°C. The absorbance of formazan solution was measured using the SpectraMax M2 plate reader at 540 nm.

### Collagen contraction assay

Collagen contraction assay was performed as previously described ^49^. Briefly, IEC were detached from the plate, counted with a T20 automated cell counter, and resuspended in DMEM medium in order to have 6,000 HT-29cf8 cells or 5,000 SK-CO15 cells per 200 μl of cell suspension. The cell suspension (200 μl) was mixed with 400 μl of collagen I solution (3 mg/ml) and 7 μl of 1M NaOH solution, and 500 μl of this mixture was added per well of a 24-well plate. After cells/collagen mixture solidified during 30 min incubation at 37 °C, 500 μl of cell culture medium was added and the gels were detached from the plastic surface by using a pipet tip. The floating collagen gels were cultured at 37 °C in 5% CO_2_ for 5 days. Images of the gels were acquired using Epson V500 photo scanner and gel area was measured using ImageJ.

### *Drosophila* UNC-45 mutants and Smurf assay

Flies were aged on standard medium for 20 days prior to performing the Smurf assay. 25 flies per vial were used during the assay. 2.5% (wt/vol) Brilliant Blue FCF dye solution (Sigma-Aldrich) was prepared in 5% sucrose solution. Small pieces of kimwipes were saturated with dye in empty vials and 20-day old flies were placed into the vials for 24 hours. 200 μl of dye was used per vial. A Smurf fly was counted when total body dye coloration was observed after 24 hours. A Leica S6 E Greenough stereomicroscope with 6.3:1 zoom was used to sort and identify Smurf flies. The average percentage of Smurf flies per vial was counted for a total of 5 vials per genotype. Statistical significance was calculated using the Mann-Whitney test.

### Statistical analysis

All data are expressed as means ± standard error (SE) from at least three biological replicates, except for FM-AFM experiments. FM-AFM data is presented as mean ± standard deviation (SD) from three independent biological replicates. Statistical analysis was performed by using a one-way ANOVA to compare multiple experimental groups (knockout with two different UNC-45A sgRNAs, or overexpression of full-length UNC-45A and different deletion mutants). A post-hoc t-test was used to compare controls with each UNC-45A-depleted group. A two-tailed unpaired Student t-test was used to compare the results obtained with two experimental groups. A Bonferroni correction was used to adjust the significance level accounting for multiple comparisons. P values < 0.05 were considered statistically significant. Statistical analysis was performed using GraphPad Prism 9.10 program.

## Results

### UNC-45A is abundantly expressed in intestinal epithelial cells and is downregulated in the colonic epithelium of ulcerative colitis patients

Immunoblotting analysis of a panel of IEC lines demonstrated abundant UNC-45A expression in well-differentiated HT-29cf8, SK-CO15, T84 and Caco-2 cells (Suppl. Fig. 1A). In HT-29cf8 and SK-CO15 cell monolayers, this cytoskeletal chaperone was enriched at the epithelial cell cortex in a close vicinity to apical junctions (Suppl. Fig. 1B,C, arrows). Furthermore, immunoblotting analysis and immunofluorescence labeling detected abundant expression of UNC-45A in normal human colonic mucosa and isolated primary human colonic epithelial cells (Suppl. Fig. 2A,B). In normal colonic mucosa UNC-45A was enriched at the apex of IEC where it colocalized with epithelial junctions (Suppl. Fig. 2A, arrows). Since previous studies reported disrupted barrier properties and altered NM II activity in the intestinal mucosa during inflammatory bowel diseases (IBD) ^50,51^, we sought to compare UNC-45A expression in patients with Crohn’s disease (CD), ulcerative colitis (UC) and non-IBD controls. Immunofluorescence labeling and immunoblotting analysis demonstrated marked decrease of UNC-45A protein level in full thickness colonic sections and isolated colonic epithelial cells of UC patients (Suppl. Fig. 2A-C). By contrast, in CD both the UNC-45A protein expression and localization appeared relatively normal (Suppl. Fig. 2A-C). Interestingly, selective down-regulation of UNC-45A in colonic epithelial cells of UC patients was accompanied by the decreased expression of NM-IIA and NM-IIC (Suppl. Fig. 2B,C).

### Depletion of UNC-45A impairs formation of the epithelial barrier and assembly of apical junctions

In order to gain insights into functional roles of UNC-45A, we used CRISPR/Cas9-mediated gene editing to down-regulate expression of this chaperon in HT-29cf8 and SK-CO15 human colonic epithelial cells. In HT-29cf8 cells, the two most efficient small guide (sg) RNAs resulted in 94-99% decrease in UNC-45A protein expression after clonal selection (Fig. 1A). In SK-CO15 cells, we selected a UNC-45A-deficient clone with ~ 97% downregulation of the targeted protein (Fig. 1B). In both IEC lines, loss of UNC-45A did not alter expression of two major IEC myosin II motors, NM-IIA and NM-IIC (Fig. 1A).

**Figure 1.**
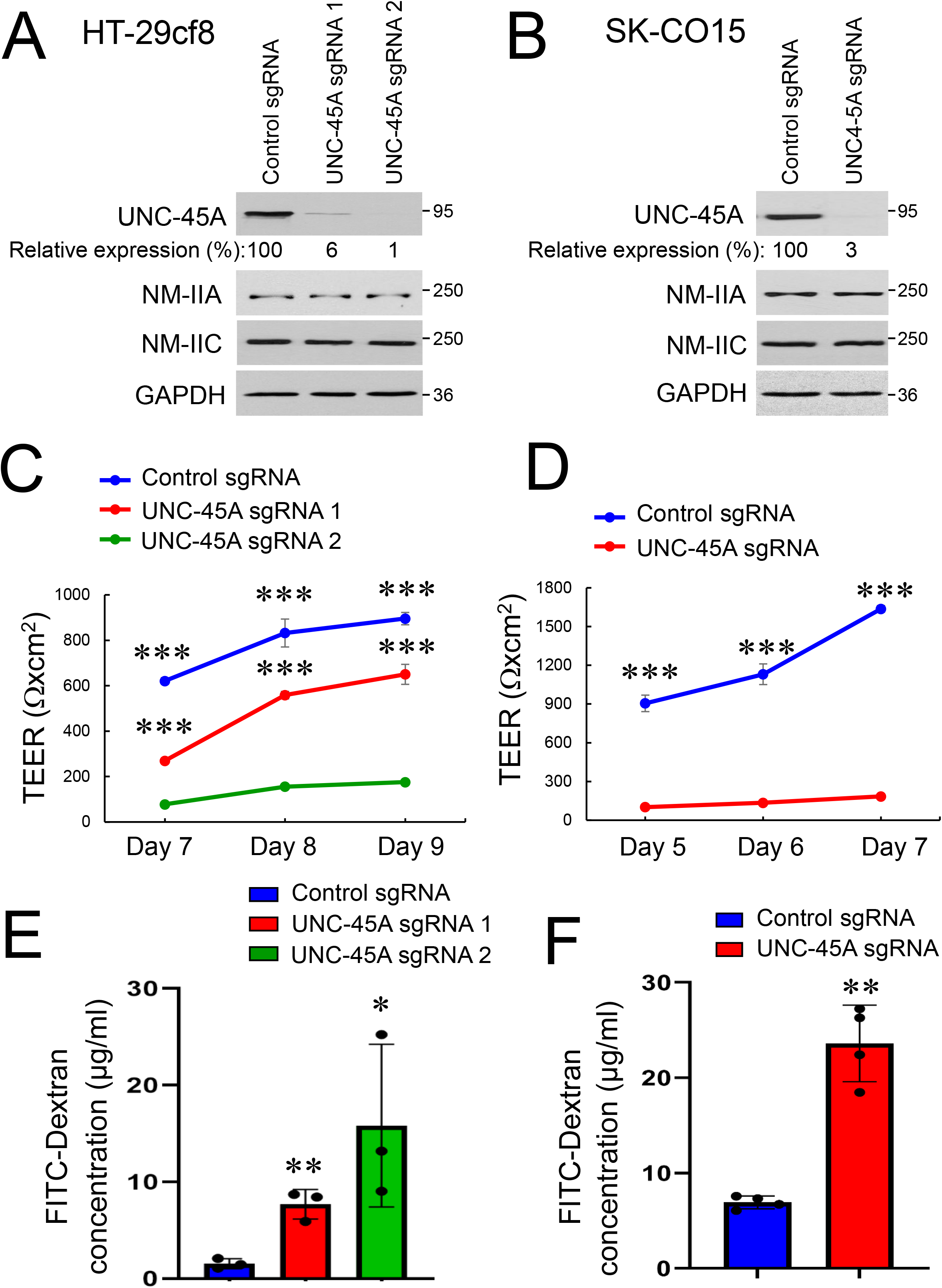
Loss of UNC-45A expression increases paracellular permeability of intestinal epithelial cell monolayers. (A,B) Immunoblotting analysis of UNC-45A and epithelial NM-II isoform expression in HT-29cf8 (A) and SK-CO15 (B) human intestinal epithelial cells with CRISPR/Cas9 mediated knockout of UNC-45A. (C,D) Transepithelial electrical resistance (TEER) of control and UNC-45A-depleted HT-29cf8 (C) and SK-CO15 (D) cell monolayers at different times post-plating. (E,F) Transmonolayer flux of FITC-dextran in control and UNC-45A-depleted IEC monolayers on day 9 (HT-29cf8, E) and day 7 (SK-CO15, F) post-plating. Data are presented as means ± SE (n=3 for HT-29cf8 and n=4 for SK-CO15 cells); *P< 0.05, **P< 0.005, ***P< 0.0005. A-F data are representative of at least three independent experiments.

Effects of UNC-45A knockout on the development of the epithelial barrier were examined by measuring transepithelial electrical resistance (TEER) and transmonolayer flux of FITC-labeled dextran. UNC-45A-depleted HT-29cf8 and SK-CO15 cells developed significantly lower TEER (Fig. 1C,D) and had higher dextran flux (Fig. 1E,F) as compared to the control cell lines, which indicates compromised barrier integrity. Such barrier dysfunction was accompanied by focal disruption of apical junctions that was observed by immunofluorescence labeling of specific markers of both AJ (E-cadherin) and TJ (ZO-1 and occludin) (Fig. 2). While in control cell monolayers AJ and TJ proteins displayed a continuous ‘chicken wire’ labeling pattern (Fig. 2A,B, arrows), this pattern was disrupted due to mislocalization of junctional proteins in UNC-45A-depleted IEC (Fig. 2, A,B, arrowheads). Interestingly, loss of UNC-45A resulted in marker alterations in cell morphology, manifested by transformation from the orthogonal to the elongated cell shape in HT-29cf8 monolayers and increased cell size in SK-CO15 monolayers (Fig. 2, A,B).

**Figure 2.**
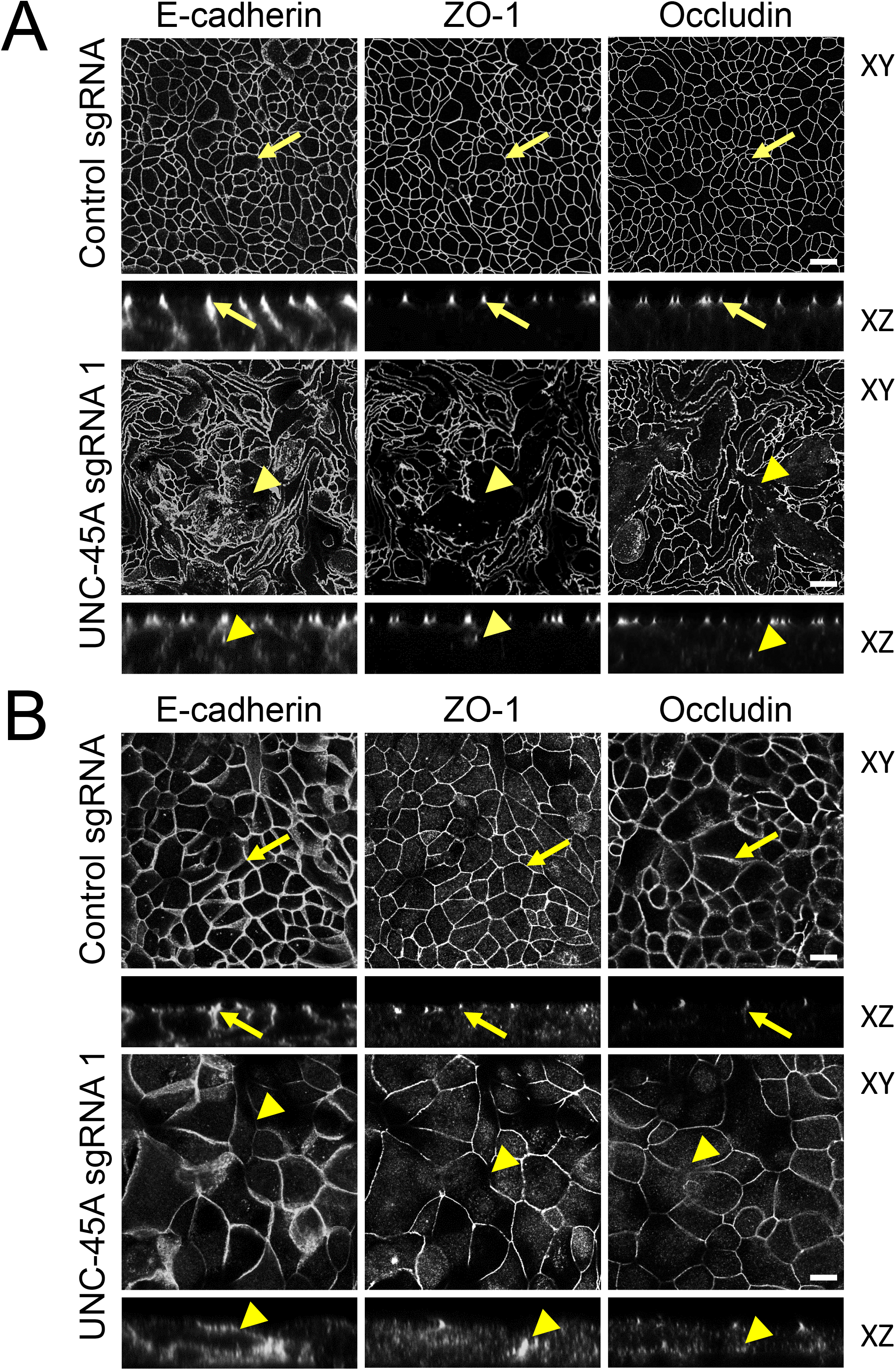
Downregulation of UNC-45A expression impairs assembly of epithelial adherens and tight junctions. Confocal microscopy images of control and UNC-45A-depleted HT-29cf8 (A) and SK-CO15 (B) cell monolayers immunolabeled for different AJ (E-cadherin) and TJ (ZO-1, occludin) proteins. Arrows indicate intact AJ and TJ in control cell monolayers. Arrowheads point to junctional disassembly in UNC-45A-depleted cells. Scale bars, 20 μm. Images shown are representative of at least three independent experiments with multiple images taken per slide.

### Loss of UNC-45A disrupts assembly of the perijunctional actomyosin cytoskeleton

Since assembly and barrier properties of epithelial junctions depend on their association with the actomyosin cytoskeleton, we rationalized that loss of UNC-45A could compromise AJ/TJ integrity by impairing assembly of the perijunctional actomyosin belt. To test this hypothesis we visualized junction-associated actin filaments and NM-II motors using fluorescence labeling and confocal microscopy. Consistent with previously published data ^35,36^, control IEC monolayers displayed a prominent perijunctional F-actin belt, enriched in both NM-IIA and NM-IIC (Fig. 3A,B, arrows). In contrast, this F-actin belt was either markedly distorted or poorly assembled in UNC-45A knockout HT-28cf8 and SK-CO15 cells, respectively (Fig. 3A,B, arrowheads). Furthermore, loss of UNC-45A significantly diminished recruitment of both NM IIA and NM IIC to the circumferential F-actin belt (Fig. 3A,B, arrowheads).

**Figure 3.**
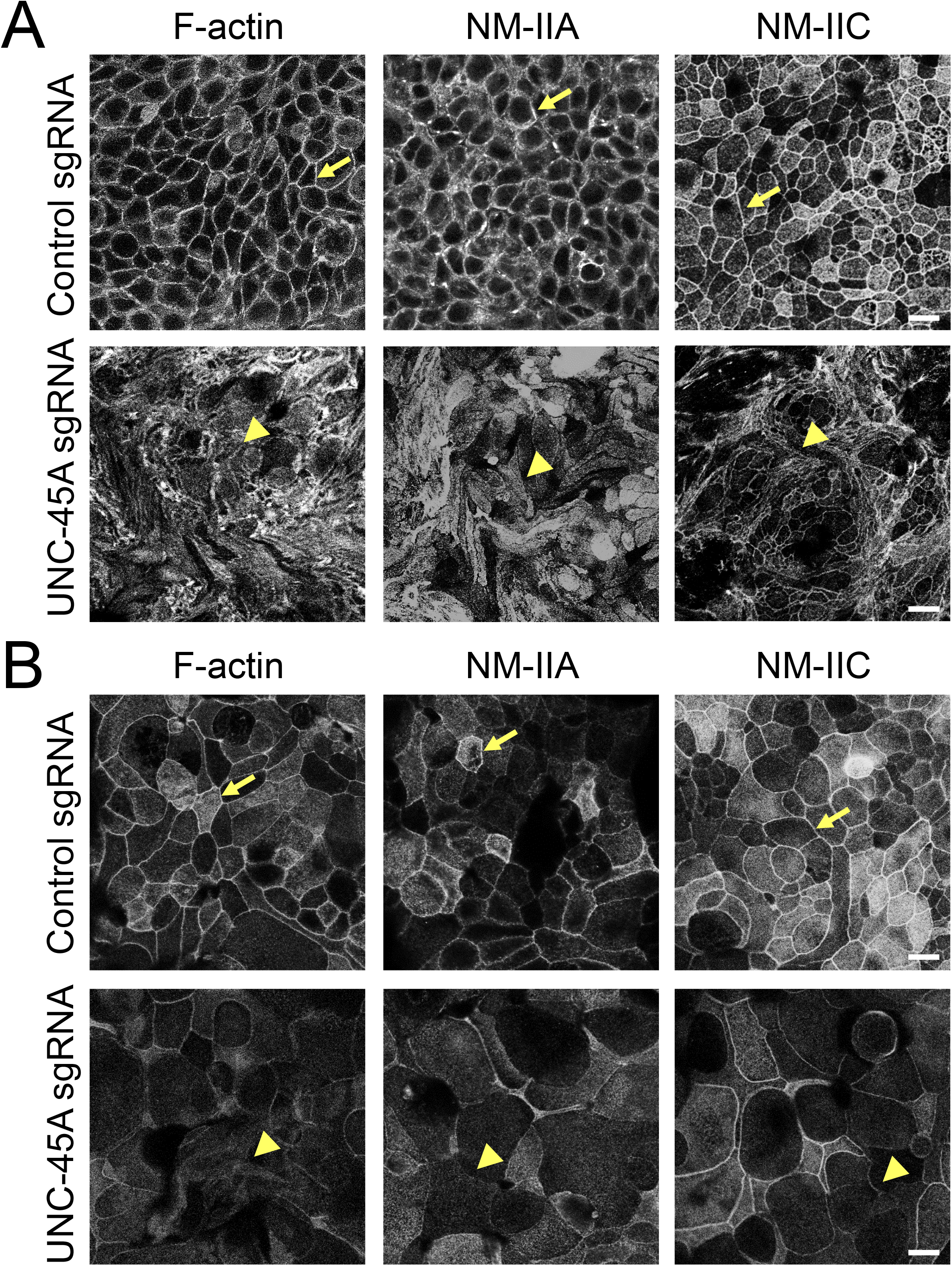
Loss of UNC-45A expression impairs organization of the perijunctional actomyosin belt in intestinal epithelial cell monolayers. Confocal microscopy images of control and UNC-45A-depleted HT-29cf8 (A) and SK-CO15 (B) cell monolayers fluorescently labeled for F-actin and major epithelial myosin II motors, NM-IIA and NM-IIC. Arrows indicate perijunctional actomyosin structures in control IEC. Arrowheads show disorganization of perijunctional NM-II bundles in UNC-45A-depleted IEC. Scale bars, 20 μm. Representative of at least three independent experiments with multiple images taken per slide.

Importantly, our data indicate physical interactions between UNC-45A and NM-IIA, which is the predominant NM-II paralog in IEC ^52,53^. NM-IIA was co-immunoprecipitated with UNC-45A from SK-CO15 cell lysates (Suppl. Fig. 3A). Furthermore, a proximity ligation assay detected extensive interactions between UNC-45A and NM-IIA in intact HT-29cf8 and SK-CO15 cell monolayers, with such interactions being greatly diminished after UNC-45A depletion (Suppl. Fig. 3B).

### Loss of UNC-45A attenuates reassembly of apical junctions and the perijunctional cytoskeleton during calcium switch

The impaired structure and permeability of the IEC barrier may not be specific consequences of UNC-45A depletion and could also reflect the delayed establishment of epithelial monolayers due to decreased cell proliferation. Indeed, loss of UNC-45A significantly delayed growth of SK-CO15, but not HT-29cf8 cells without causing cell apoptosis (Suppl. Fig. 4A-D). To establish the specific effects of UNC-45A depletion on AJ/TJ assembly, we used a classical calcium switch model in confluent IEC monolayers that enables selective examination of junctional dynamics ^35–37^. The described experiments were performed in SK-CO15 cells, because HT-29cf8 cells did not readily restore TJ integrity in this model. Re-addition of extracellular calcium caused rapid (within 5 h) AJ and TJ reassembly in control SK-CO15 cell monolayers (Fig. 4A arrows, B). In contrast, restoration of apical junctions was significantly delayed in UNC-45A-deficient cells (Fig. 4A, arrowheads, B). Calcium driven AJ/TJ reassembly in control IEC was accompanied by reformation of the sharp, NM-IIA and NM-IIC-containing circumferential actomyosin belt (Fig. 4C arrows, D). The assembly of this belt was inhibited in UNC-45A deficient IEC that displayed a loose, non-condensed arrays of parallel myosin bundles at the intercellular contacts (Fig. 4C arrowheads, D).

**Figure 4.**
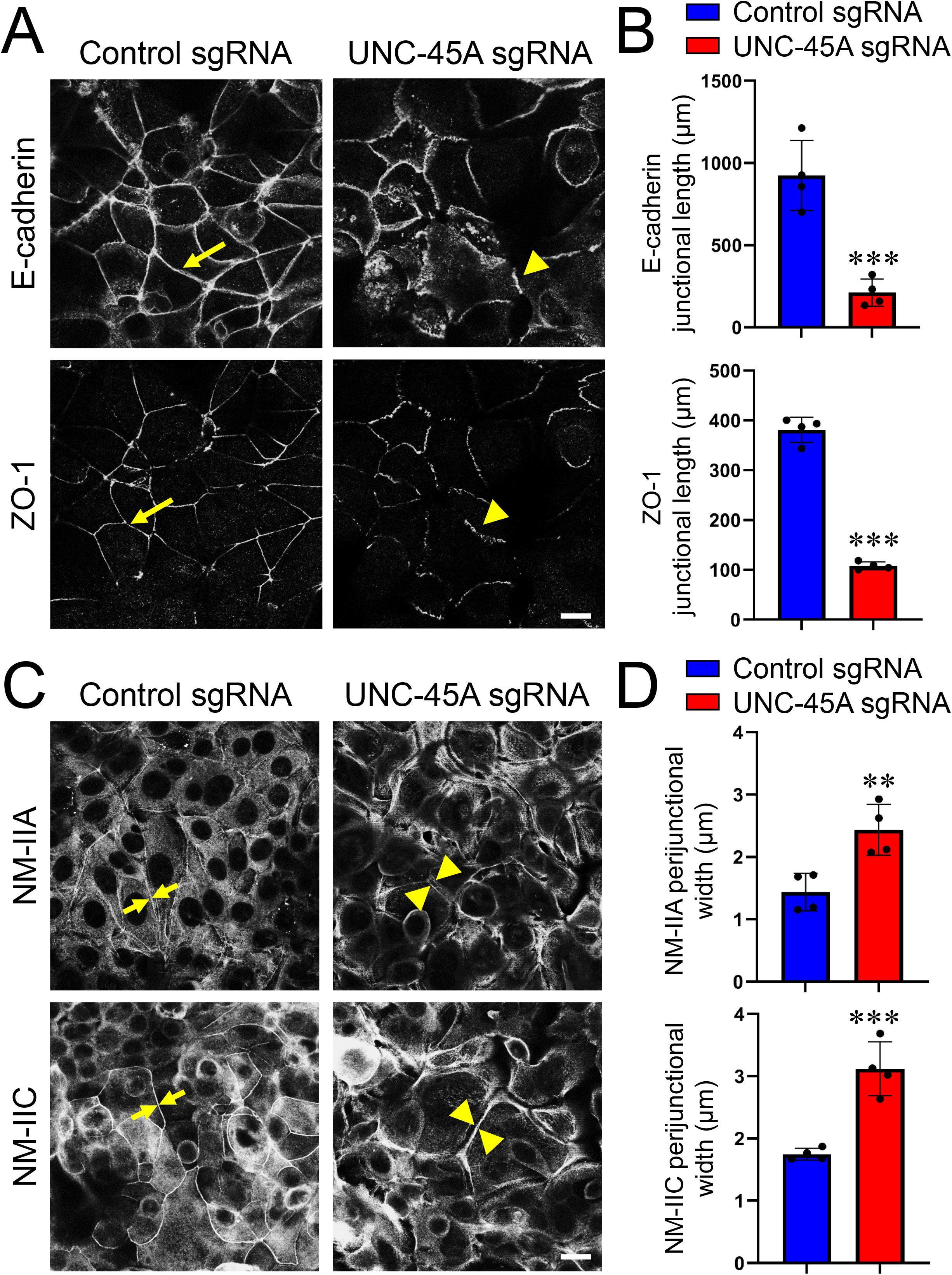
Depletion of UNC-45A attenuates AJ/TJ reassembly and reformation of the perijunctional myosin belt during the extracellular calcium switch. (A,B) Immunolabeling of E-cadherin and ZO-1 in control and UNC-45A knockout SK-CO15 cells after 5 h of extracellular calcium repletion. Representative confocal microscopy images (A) and quantification of junctional length (B) is shown. Each dot represents an averaged junctional length measured in different images. (C,D) Immunolabeling of NM-IIA and NM-IIC in control and UNC-45A knockout SK-CO15 cells after 5 h of extracellular calcium repletion. Representative confocal microscopy images (C) and quantification of the width of the perijunctional NM-II belt (D) is shown. Arrows indicate AJ/TJ reassembly and restoration of the actomyosin belt in control IEC. Arrowheads show poorly assembles apical junctions and perijunctional NM-II bundles in UNC-45A knockout IEC. Each dot represents an averaged width of 200 cell-cell contacts in four different images. Means ± SE (n=4); **P< 0.005, ***P< 0.0005. Scale bar 20 μm. A-D data are representative of three independent experiments.

### Loss of UNC-45A inhibits IEC migration

Assembly and dynamics of apical junctions plays critical roles not only in regulating epithelial barriers, but also in restoring integrity of injured epithelial cell monolayers, by controlling collective cell migration ^2,3^. Given the observed role of UNC-45A in regulating junctional integrity, we sought to examine if this chaperone also influences on IEC motility. Loss of UNC-45A significantly delayed collective migration on HT-29cf8 and SK-CO15 cell monolayers in a wound healing assay (Fig. 5A-D). Furthermore, depletion of this chaperone caused a marked inhibition of transfilter migration of individual IEC in the Boyden Chamber (Fig. 5E-H). The latter result uncouples the effects of UNC-45A inhibition on epithelial junctional integrity and cell migration, and suggests its independent functions in stationary and motile IEC. To gain insights into the mechanism of UNC-45A regulation of IEC migration, we examined whether it controls cell adhesion to the extracellular matrix (ECM), which represents a key step of cell motility ^39,54^. UNC-45A depletion significantly increased adhesion of HT-29cf8 and SK-CO15 to collagen I (Fig. 5I-L). Interestingly, the hyperadhesiveness of UNC-45A-deficient IEC was accompanied by the increased number of focal adhesions that mediate cell-ECM interactions and decreased abundance of actomyosin bundles at the migrating cell edge (Suppl. Fig. 5).

**Figure 5:**
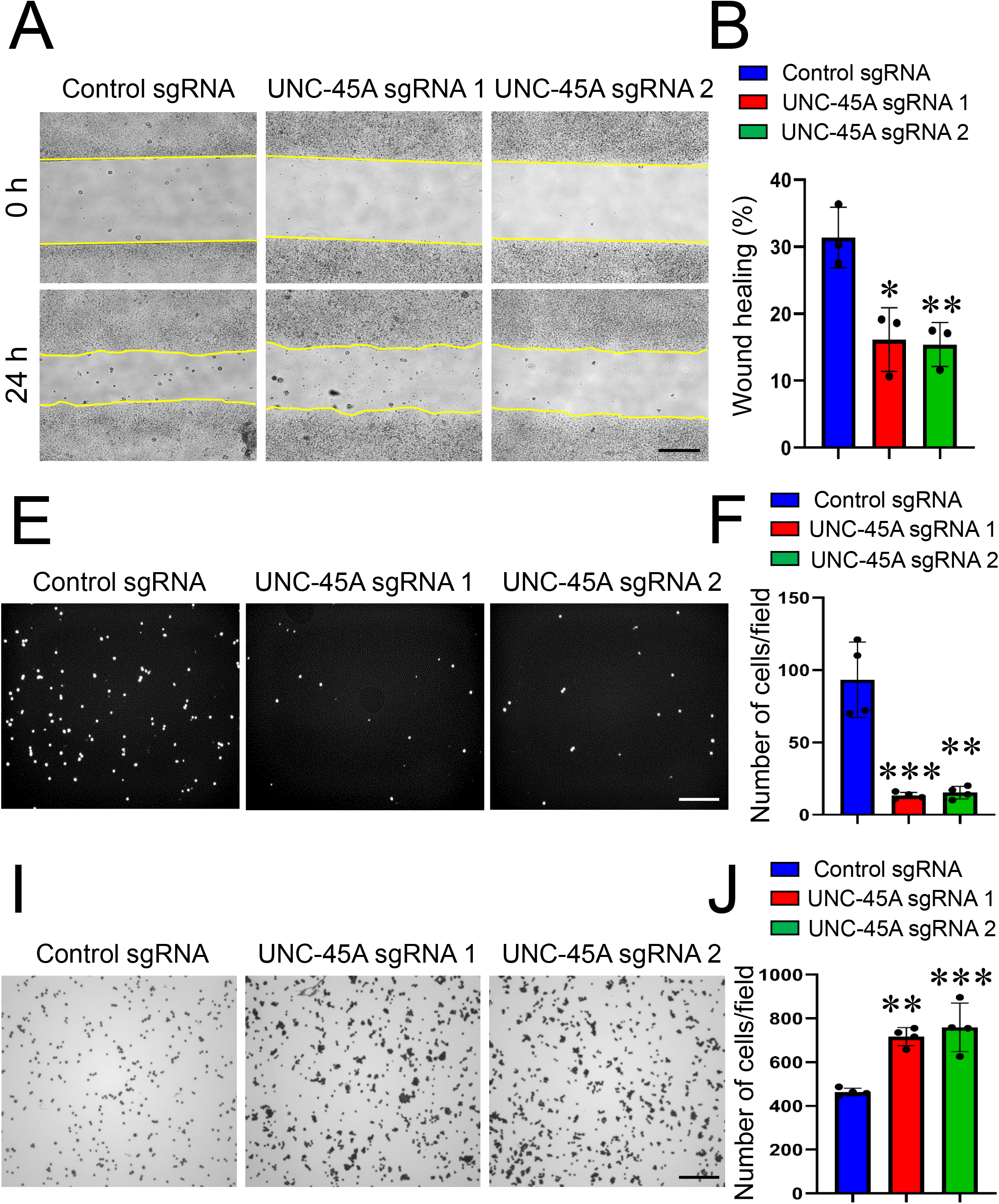
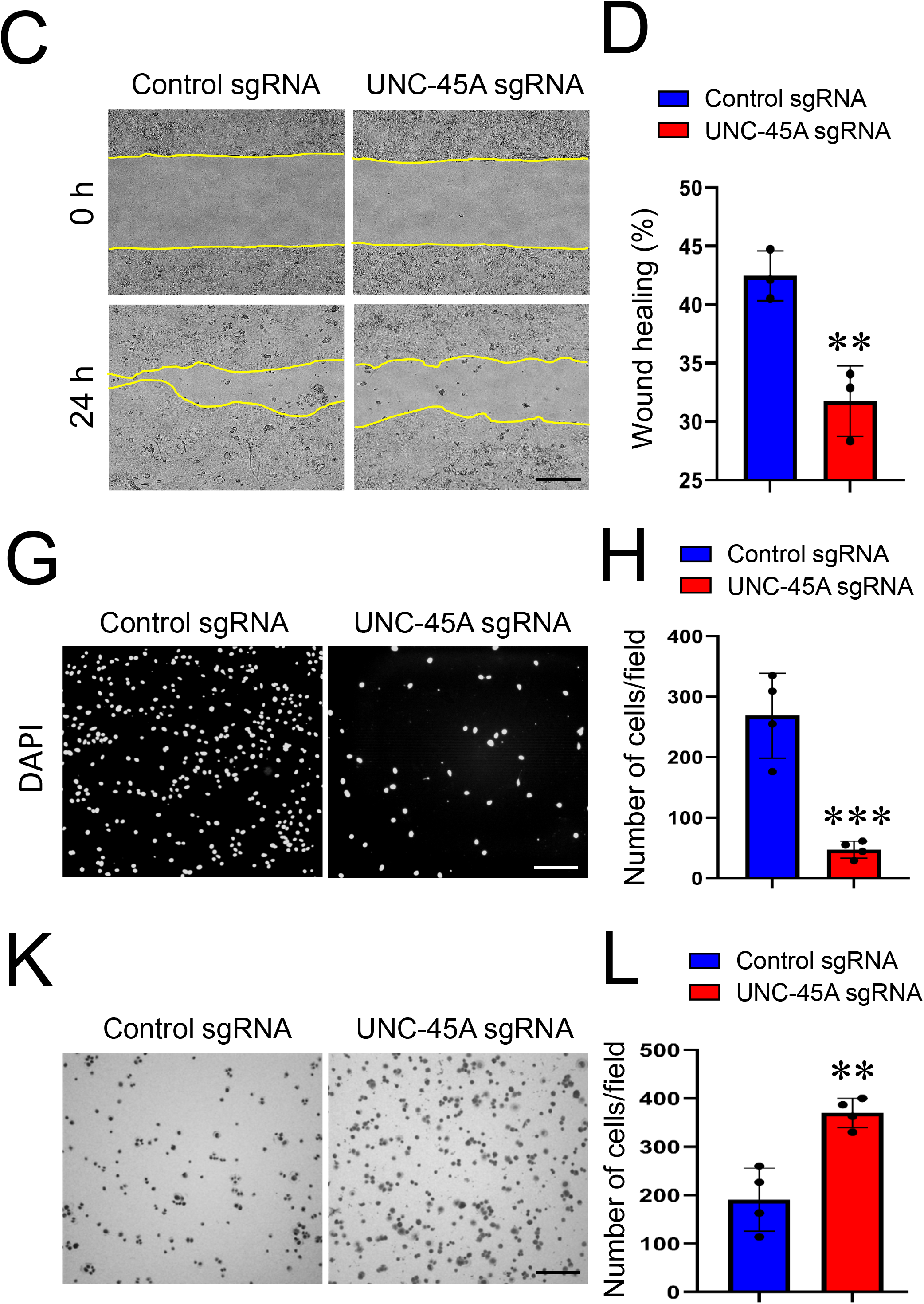
Loss of UNC-45A expression inhibits epithelial cell migration and increases cell ECM adhesion. (A-D) Wound healing assay in control and UNC-45A-depleted HT-29cf8 (A,B) and SK-CO15 (C,D) cell monolayers. Representative wound images (A,C) and quantification of wound closure (B,D) are shown. (E-H) Boyden Chamber migration assay of control and UNC45A-knockout HT-29cf8 (E,F) and SK-CO15 (G,H) cells. Representative images of migrated cells (E,G) and quantification of transfilter cell migration (F,H) are shown. (I-L) Collagen I matrix adhesion assay of control and UNC45A-knockout HT-29cf8 (I,J) and SK-CO15 (K,L) cells. Representative images of migrated cells (I,K) and quantification of cells adhered after 30 min incubation (J,L) are shown. Mean ± SE (n =3); *P< 0.05, **P< 0.005; ***P< 0.0005, as compared to the control sgRNA group. Scale bars, 200 μm. A-D data are representative of at least three independent experiments, E-K data are representative of two experiments.

### Loss of UNC-45A inhibits IEC supracellular tension and contractility

Our data indicate that UNC-45A controls both IEC apical junctions and cell-matrix adhesions by regulating proper assembly of actomyosin structures associated with different adhesion complexes. Since NM-II is known to strengthen epithelial junctions and focal adhesions by generating mechanical forces, we hypothesized that loss of UNC-45A would diminish mechanical forces in IEC. We tested this hypothesis by examining cytoskeletal forces applied to cell-cell and cell-matrix adhesions in control and UNC-45A-deficient IEC. Three different approaches were used to examine forces: Frequency modulated acoustic atomic force microscopy (FM-AFM) was used to examine force generation at the apical IEC surface ^48^. Mechanical forces at AJ were measured by using a conformational sensitive α18 anti-α-catenin monoclonal antibody ^55,56^ and E-cadherin based mechanosensitive FRET sensor ^44,57^ Forces generated at ECM adhesions were examined using a collagen contraction assay.

Loss of UNC-45A significantly decreased the epithelial tension and viscosity of the apical surface of HT-29cf8 cell monolayers as measured by FM-AFM (Fig. 6 A,B), thereby suggesting decreased contractile forces. Furthermore, downregulation of this cytoskeletal chaperone decreased junctional labeling intensity of the α18 antibody that recognizes α-catenin conformation unfolded by the pulling mechanical forces ^56^, which also indicates decreased forces at cell-cell contacts of UNC-45A-depleted IEC (Fig. 6C,D). By contrast, expression of E-cadherin based tension sensor did not show significant effects of UNC-45A depletion on forces applied to E-cadherin bases cell-cell contacts (Suppl. Fig. 6). Finally, loss of UNC-45A significantly inhibited collagen gel contraction by embedded HT-29cf8 (Fig. 6 E,F) and SK-CO15 (data not shown), which indicates diminished actomyosin contractile forces at cell-ECM contacts. Together, these data suggest that UNC-45A is required for generation of NM-II-driven contractile forces at different adhesive structures in IEC monolayers.

**Figure 6.**
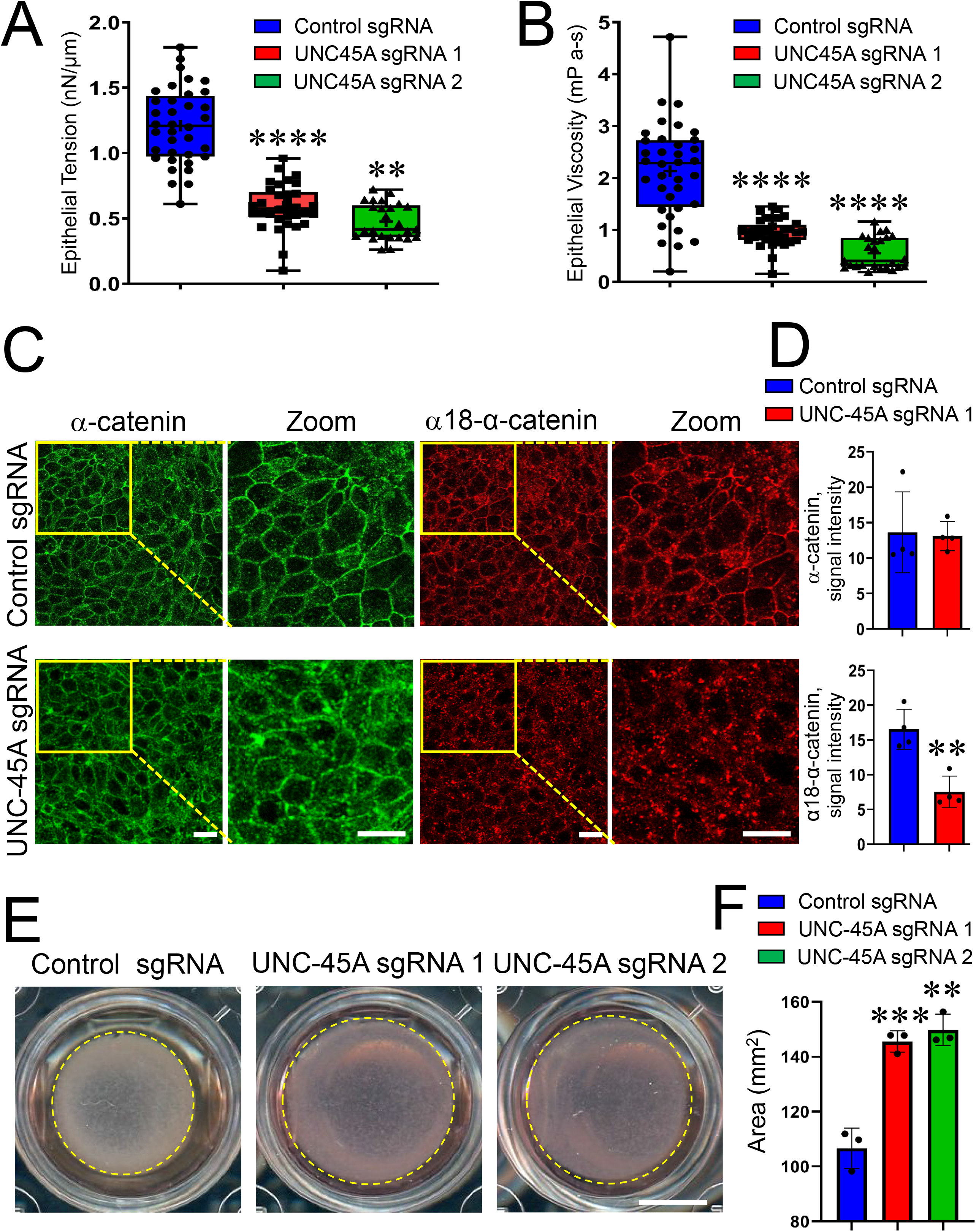
UNC-45A regulates mechanical forces in intestinal epithelial cells. (A,B) Viscoelastic properties, epithelial tension (A) and viscosity (B) measurements in control and UNC-45A-depleted HT-29cf8 cell monolayers by noncontact frequency modulation atomic force microscopy (FM-AFM). Means ± SD; number of cells for control sgRNA n=35, UNC-45A sgRNA1 n=31, and UNC-45A sgRNA2 n=26; **P< 0.005; ****P< 0.0001, as compared to the control sgRNA group. (C,D) Immunofluorescence labeling of control and UNC-45A-depleted HT-29cf8 cell monolayers with antibodies recognizing either total α-catenin, or an open conformation of α-catenin (α18-α-catenin antibody). Representative confocal images (C) and quantification of the labeling intensity (D) is shown. Each dot represents an averaged signal intensity of 200 cell-cell contacts measured in four different images. Means ± SE (n=4); **P< 0.005, as compared to the control sgRNA group. Scale bar, 20 μm. (E,F) Contraction of collaged I gel with embedded control or UNC-45A-depleted HT-29cf8 cells. Representative images of contracted gels (E) and quantification of the gel area after 4 days of contraction (F) are shown. Scale bar, 50 mm. Means ± SE (n=4); **P< 0.005; ***P< 0.0005, as compared to the control sgRNA group. A-F data are representative of three independent experiments.

### The myosin-binding domain of UNC-45A selectively perturbs barrier properties and migration of IEC monolayers

While conventional myosins are the most well-known folding clients for UNC-45A, this chaperone has additional myosin independent activities, such as interacting with tubulin and modulating microtubule dynamics ^23–25^. Since altered dynamics of microtubules could affect epithelial barrier integrity and cell migration, we next sought to examine whether this mechanism contributes to the abnormal junctional assembly and wound healing caused by UNC-45A knockout. First, we used immunofluorescence labeling to visualize microtubules, but did not observe significant alterations in the apical microtubule meshwork following UNC-45A depletion in SK-CO15 and HT-29cf8 cells (Suppl. Fig. 7A,B). Next, we created SK-CO15 cell lines stably expressing either full length UNC-45A (UNC-45A-GFP), its N-terminal truncated ΔTPR-GFP (residues 147-971), or C-terminally truncated ΔUCS-GFP (residues 1-539) (Fig. 7A). Cells expressing only GFP served as a control. Epithelial permeability measurements revealed that the ΔTPR mutant that binds to NM-II, but not to microtubules, markedly diminished TEER and increased FITC-dextran flux (Fig. 7B,C). By contrast, the ΔUCS mutant that interacts with microtubules, but lacks the myosin-folding activity modestly attenuated TEER development without affecting the FITC-dextran flux (Fig. 7B,C). Furthermore, the calcium switch assay demonstrated that expression of the ΔTPR, but not the ΔUCS mutant attenuated AJ and TJ reassembly (Fig 7. D,E). In wound healing experiments, expression of the ΔTPR UNC-45A mutant significantly inhibited collective migration SK-CO15 cells, whereas expression of the ΔUCS mutant had a slight inhibitory effect that did reach significant difference (Suppl. Fig. 8A,B). Together, these data suggest that UNC-45A regulates IEC barrier, apical junction assembly and cell migration primarily by targeting NM-II in microtubule-independent fashion.

**Figure 7.**
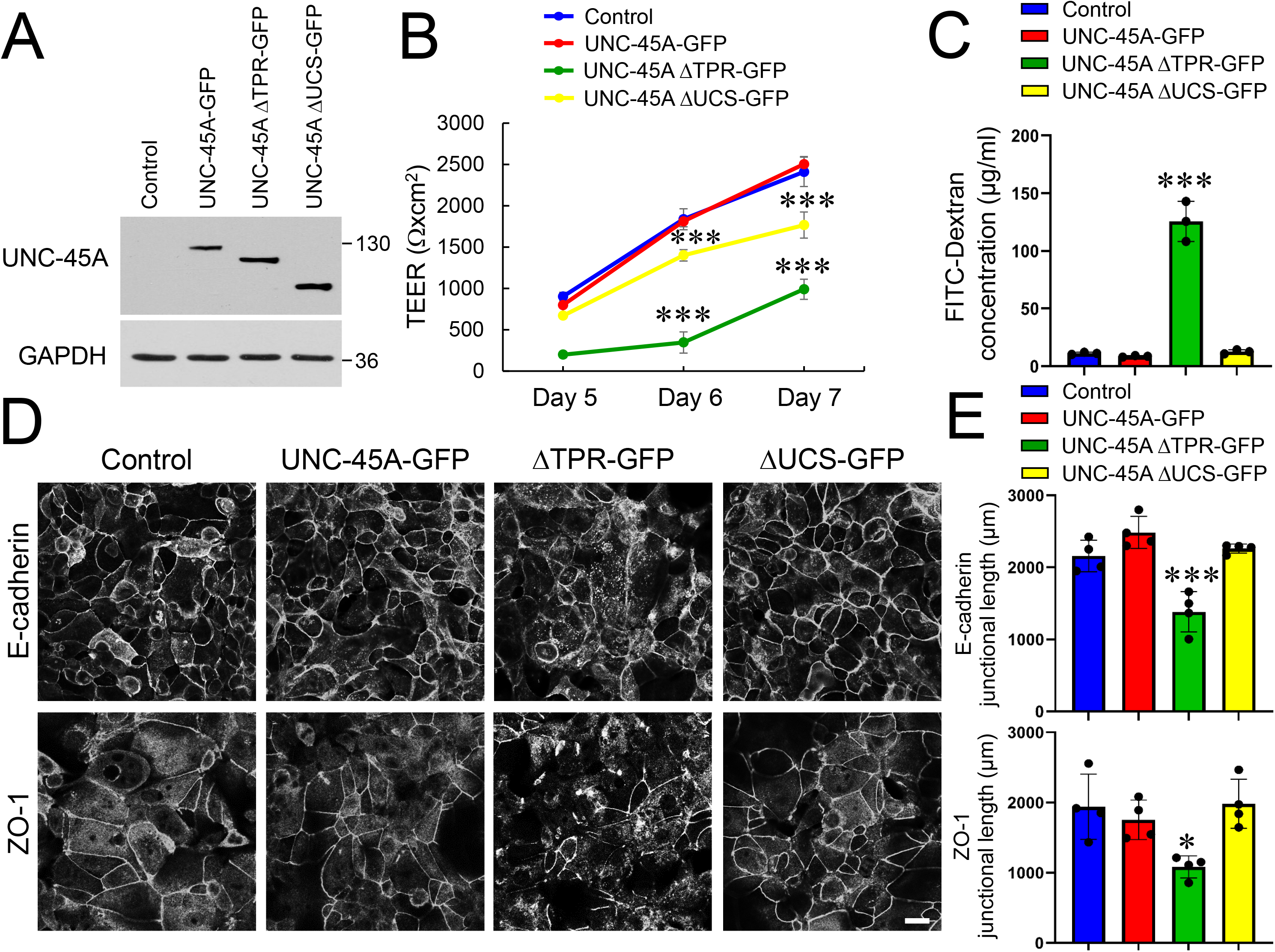
Expression of N-terminal domain deletion mutant UNC-45A disrupts epithelial barrier integrity and attenuates AJ and TJ assembly. (A) Immunoblotting analysis of SK-CO15 expressing either GFP-tagged full-length UNC-45A or two different deletion mutants of UNC-45A, ΔTPR-GFP (N-terminal truncated), or ΔUCS-GFP (C-terminally truncated). (B) Transepithelial electrical resistance (TEER) of control SK-CO15 cell monolayers and cells expressing UNC-45A constructs at different times post-plating. (C) Transmonolayer flux of FITC-dextran in control SK-CO15 cell monolayers and cells expressing different UNC-45A constructs on day 7 post-plating. (D,E) Immunolabeling of E-cadherin and ZO-1 in control SK-CO15 cell monolayers and cells expressing different UNC-45A constructs after 5 h of extracellular calcium repletion. Representative confocal microscopy images (D) and quantification of junctional length (E) is shown. Means ± SE (n=3 for B,C; n=4 for D); *P< 0.05, ***P< 0.0005. Scale bar, 20 μm. A-E data are representative of three independent experiments.

### UNC-45 regulates permeability of *Drosophila* gut

In order to validate the *in vivo* relevance of our finding in IEC monolayers we sought to determine the effects of lowering UNC-45 expression on permeability of the *Drosophila* gut. Two mutant fly strains with monoallelic depletion of UNC-45 were examined since homozygous deletion of this chaperone results in *Drosophila* lethality ^31^. Immunoblotting analysis demonstrated approximately 30-50% reduction of UNC-45 expression in tissues of mutant flies (Fig. 8A). Aged flies were fed with a sucrose solution containing Brilliant Blue dye, and then scored as to whether their intestines were leaky, based on the whether or not dye had leaked into extra-intestinal tissues (‘Smurf” phenotype) (Fig. 8B) ^58,59^. Increased numbers of Smurf flies were observed in both 20 day-old females and males of the unc-45^03692/+^ mutant genotype characterized by 50% UNC-45 depletion (Fig. 8C). Another mutant genotype with less pronounced UNC-45 downregulation, unc-45^EY03043-33/+^, showed a tendency toward increased gut permeability, however numbers of Smurf flies did not reach statistical significance (Fig. 8C). These data suggest that UNC-45 expression could serve as a ‘rheostat’ controlling permeability of the *Drosophila* gut.

**Figure 8.**
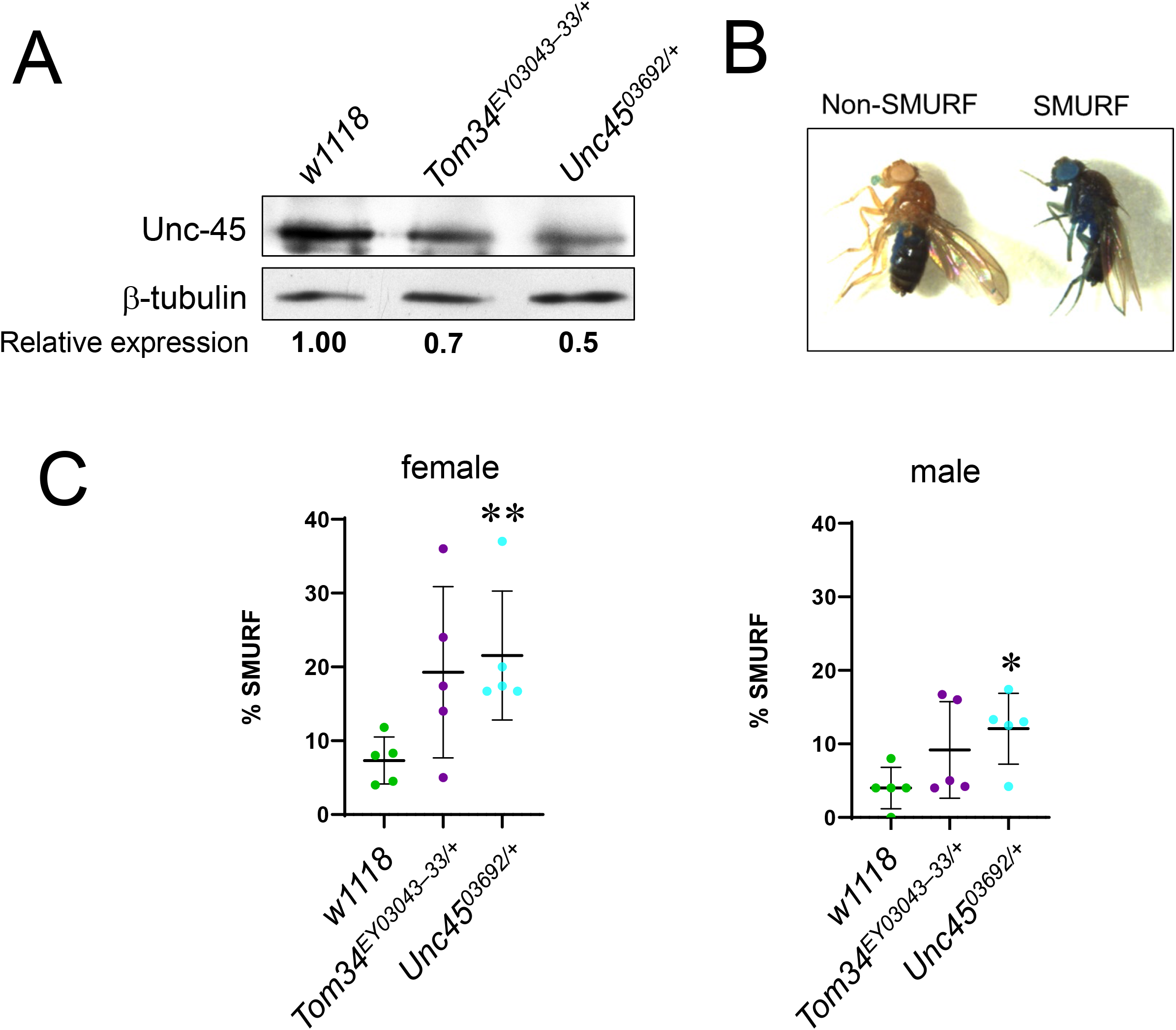
Reduced expression of Unc-45 in *Drosophila* increases intestinal permeability. (A) Immunoblotting analysis of Unc-45 protein in wild type (*w1118*) or heterozygous unc-45 mutant (unc-45^EY03043-33^ or unc-45^03692^) *Drosophila melanogaster* adult flies. β-tubulin was used as a loading control. (B) Pictures of adult *Drosophila* females categorized as non-Smurf (blue dye localized to intestine and proboscis) or Smurf (blue dye spread from intestine to the majority of tissues). (C) Quantitation of the percentage of flies exhibiting the Smurf phenotype (right picture in B) for each gender of each genotype analyzed in A. Each dot represents an individual experiment containing 25 flies. *p<0.05, **p<0.01.

## Discussion

NM-II is a crucial cytoskeletal motor that regulates assembly and functions of epithelial apical junctions and basal ECM adhesions ^12,60,61^. While molecular mechanisms that govern NM-II ATPase activity and its dynamic interactions with actin filaments have been extensively investigated, much less is known about the regulation of NM-II organization into higher-order contractile structures associated with epithelial adhesive complexes. In this study we describe a novel mechanism controlling NM-II-dependent assembly of apical junctions and ECM adhesions that is essential for the integrity and repair of the intestinal epithelial barrier. This mechanism involves a cytoplasmic myosin chaperone, UNC-45A. Although an essential role for UNC-45A in controlling myosin folding and function was originally recognized more than two decades ago ^62^, its involvement in the regulation of intercellular adhesions and tissue barriers has not been previously investigated.

Our data suggest that UNC-45A regulates two major permeability pathways in IEC monolayers. One is a paracellular pore pathway selectively permeable to small ions, and the other is a leak pathway permeated by large molecules ^4^. Disruption of the pore and the leak pathways in UNC-45A deficient IEC cells was revealed by decreased TEER and increased FITC-dextran flux, respectively (Fig. 1). The described defects in barrier permeability were associated with disruption of AJ and TJ integrity (Fig. 2). Importantly, loss of UNC-45A markedly attenuated epithelial junctional reassembly during extracellular calcium switch (Fig. 4), which signifies specific regulatory roles of this chaperone at IEC apical junctions.

Several lines of evidence suggest that loss of UNC-45A disrupts the integrity and remodeling of epithelial junctions by impairing assembly and contractility of the perijunctional actomyosin cytoskeleton. Thus, UNC-45A physically interacts with NM-IIA in IEC (Suppl. Fig. 3), and its loss markedly disrupts organization of the circumferential actomyosin belt associated with steady-state and reassembling junctions (Figs. 3,4). This phenocopies barrier disruption and attenuated AJ/TJ reassembly that was previously described in human epithelial cells after either pharmacologic inhibition of NM-II motors, or genetic depletion of NM IIA ^35,36,63^. Since the sarcomeric organization of NM-II filaments within the perijunctional belt resembles organization of muscles myofibrils ^10,13^, the observed disorganization of the perijunctional actomyosin belt in UNC-45A-deficient IEC resembles impaired myofibril assembly caused by loss of this chaperone in cultured muscle cells and model organisms ^16,18,31^.

Consistent with disordered organization of the perijunctional actomyosin cytoskeleton, loss of UNC-45A reduced myosin-dependent forces at epithelial junctions. This was manifested by decreased epithelial tension and viscosity of the apical surface of UNC-45A-depleted IEC (Fig. 6A,B), which was previously shown to be dependent by myosin contractility ^48^. Since the applied atomic force microscopy technique measures mechanical properties of epithelial monolayers with depth up to 2.5 μm, our experiments most likely detected decreased mechanical forces applied to both TJ and AJ in UNC-45A-depleted cell. This is consistent with the results obtained using a conformation-sensitive α-catenin antibody that also detected inhibition of pooling forces at AJ/lateral cell-cell contacts caused by UNC-45A-deficient IEC (Fig. 6C,D). Surprisingly, no changes in mechanical forces have been observed by using FRET-based E-cadherin tension sensor (Suppl. Fig. 6). It is important to consider that the E-cadherin tension sensor measures average force per E-cadherin molecule, contrasting to the applied atomic force microscopy technique, which measures the overall mechanical force state between cells. Additionally, UNC-45A deletion likely affects the mechanical forces across other load-bearing cell adhesion proteins such as TJs which have been recently shown to be subject to actomyosin-driven forces ^64,65^. Furthermore, the conflicting data obtained with the E-cadherin tension sensor and the conformation-sensitive α-catenin antibody could be due to α-catenin interactions with other types of cadherin (e.g., P-cadherin, LE-cadherin, etc.) that are abundantly expressed in IEC. It is possible that loss of UNC-45A inhibits forces at specific cadherin-catenin domains of IEC junctions.

Finally, expression of UNC-45A deletion mutants indicates that this chaperone regulates epithelial barrier and AJ/TJ assembly in an NM-II-dependent fashion. Indeed, we found that the N-terminal deletion mutant that retains the myosin-binding domain (ΔTPR-UNC-45A) markedly disrupts barrier integrity, attenuates AJ and TJ reassembly and inhibits collective cell migration (Fig. 7, Suppl. Fig. 8). Consistently, a previous study demonstrated that expression of this mutant resulted in the assembly of abnormal stress fibers in osteosarcoma cells ^30^. What are the mechanisms by which the ΔTPR UNC-45A mutant disrupts the IEC barrier? While the C-terminal UCS domain is sufficient to fold myosin in vitro ^19^, the entire UNC-45A structure containing both the UCS and TPR domains is required to build the sarcomeric organization of muscle myofibrils. Both domains are involved in intermolecular interactions that mediate UNC-45 oligomerization into periodic tandem modules thereby providing a template for orchestrated self-assembly of multiple NM-II molecules ^20^. Therefore, the exogenous ΔTPR mutant is likely to compete for NM-II binding with the endogenous full-length chaperone, preventing assembly of high order myosin filaments. By contrast, the mutant overexpression experiments did not find strong evidence that interactions with microtubules contribute to UNC-45A-dependent regulation of IEC barrier integrity and junctional reassembly (Fig. 7; Suppl. Figs. 7 & 8). This is consistent with our previous study indicating that microtubules are dispensable for apical junction assembly in IEC ^66^ and further signifies NM-II-dependent regulation of epithelial barrier integrity by UNC-45A.

UNC-45A appears to play dual functional roles in IEC by both regulating barrier integrity, and regulating cell migration. Generally, the roles of UNC-45A in mammalian cell migration remain controversial. Depletion of this chaperone was shown to inhibit motility of breast and ovarian cancer cells ^27,29^, but promoted migration of osteosarcoma and melanoma cells ^30,67^. This resembles the reported opposite effects of NM-II inhibition on cell migration under different experimental conditions ^52,68^ and may reflect complex regulatory effects of UNC-45A-driven myofilament assembly in motile cells. We observed that loss of UNC-45A inhibited both collective migration of IEC sheets and individual cell passage through membrane pores (Fig. 5A-H). Strikingly, the attenuated collective migration of UNC-45A-depleted IEC was coupled with increased cell adhesion to ECM and elevated number of FA (Fig. 5I-L; Suppl. Fig. 5A,B). Furthermore, hyperadhesiveness of UNC-45A-depleted IEC was accompanied by decreased contractile forces at the ECM adhesions and disorganization of F-actin bundles at the migrating cell edge (Fig. 6E-F; Suppl. Fig. 5C). Decreased myosin-driven forces are likely to prevent efficient FA remodeling thereby attenuating the orchestrated directional movement of IEC monolayers. Likewise, inhibited NM-II contractility could impair cell deformability, which is required for efficient migration through confined spaces, such as membrane pores. This could explain attenuated transfilter migration of UNC-45A-deficient IEC. Similar to the effects on the IEC barrier, UNC-45A regulation of IEC migration appears to be dependent on its interactions with myosin motors, but not microtubules. This conclusion is supported by the unique dominant negative effect of exogenously expressed myosin-binding mutant of UNC-45A (Suppl. Fig. 8). Together, our results uncover roles for UNC-45A in controlling multiple functions of NM-II in the intestinal epithelium.

*Drosophila* UNC-45 is an essential protein that is expressed at all developmental stages. Like human UNC-45A, *Drosophila* UNC-45 also colocalizes with non-muscle myosin II ^31^. Furthermore, unc-45 mutant embryos display defective myosin accumulation in body wall muscles at 22 hours post-fertilization ^31^. Our results demonstrate that the intestines of 20-day old unc-45 mutant adults display increased permeability, as compared to wild-type controls (Fig. 8). Although the functions of UNC-45 in myosin regulation have not been investigated in the adult *Drosophila* gut, it is possible that a conserved role for UNC-45 in promoting myosin assembly at TJ and AJ exists in the *Drosophila* gut, and this contributes to the integrity of the adult *Drosophila* intestinal epithelial barrier.

In addition to characterizing novel roles of UNC-45A in regulating barrier integrity and repair in IEC monolayers, our study suggests possible dysfunctions of this molecular chaperone during mucosal inflammation. Specifically, we found marked and selective downregulation of UNC-45A protein expression in the colonic epithelium of UC patients (Suppl. Fig. 2). Interestingly, expression of both NM-IIA and NM-IIC was also diminished in IEC isolated from UC patients (Suppl. Fig. 2B,C), although we do not know whether loss of UNC-45A plays causal role in the depletion of NM-II motors in the inflamed intestinal mucosa. This chaperone was found to be upregulated during tumorigenesis ^27,29^ and we present the first evidence for its altered expression under inflammatory conditions. While we did not investigate molecular mechanisms of the disease-associated downregulation of UNC-45A, its expression is known to be tightly controlled in muscle cells via the ubiquitin-proteosomal degradation system ^29,69^. Interestingly, a loss-of-function mutation in UNC-45A in humans resulted in chronic diarrhea, among other clinical symptoms ^70^. This suggests that UNC-45A function is essential for normal intestinal epithelial homeostasis, and the observed decrease in its expression during mucosal inflammation in UC patients could impair cytoskeleton-dependent processes in IEC, including establishment of paracellular barrier and healing of mucosal wounds.

In conclusion, our study identified a novel mechanism that control actomyosin-dependent integrity and remodeling of the intestinal epithelial barrier. This mechanism involves a myosin chaperone, UNC-45A. UNC-45A binds to NM-IIA in cultured IEC and controls NM-II assembly and contractility at epithelial adhesions. Through its myosin folding activity, this molecular chaperone plays a dual role in epithelial monolayers by regulating formation of apical junctions and collective cell migration during wound healing. Loss of UNC-45A expression during intestinal inflammation may promote leakiness of the gut barrier and impede mucosal restitution.

## Acknowledgments

We thank Dr. Nayden Naydenov for help with isolating primary human colonic epithelial cells. This work was supported by National Institute of Health grants RO1 DK108278 and RO1 DK126702 to A.I.I, R35 GM119617 to D.E.C, R21 AI153780 to M.S.L. and by funding from Cure for IBD and a Velosano Award to M.S.L. This work was also supported by the National Institutes of Health (NIH) Intramural Research Program of the National Institutes of Biomedical Imaging and Bioengineering (grant ZIA EB000094) and the NIH Distinguished Scholars Program to A.X.C.R. Confocal microscopy was performed at the Lerner Research Institute Digital Imaging Microscopy Core.

## Supplemental Figures

**Supplemental Figure 1. Expression and localization of UNC-45A in polarized model intestinal epithelial cell monolayers.**

(A) Immunoblotting analysis of UNC-45A expression in a panel of different human intestinal epithelial cell lines.

(B,C) Dual immunofluorescence labeling of UNC-45A (green) and β-catenin (red) in polarized HT-29cf8 (B) and SK-CO15 (C) cell monolayers. Representative microscopic images of apical (xy) and lateral (xz) sections are shown. Arrows indicate enrichment of UNC-45A at apical junctions. Scale bars, 20 μm. A and B/C data are representative of two and three independent experiments, respectively.

**Supplemental Figure 2. UNC-45A expression is down-regulated in the human colonic epithelium of patients with ulcerative colitis.**

(A) Dual-immunofluorescence labeling of UNC-45A (red) and E-cadherin (green) in whole thickness sections of normal colonic mucosa and colonic tissue of a Crohn’s disease or ulcerative colitis patient. Arrows indicate UNC-45A enrichment at apical junctions in normal mucosa, and arrowheads show the loss of UNC-45A in ulcerative colitis mucosa. Images are representative of 6 patients per group.

(B,C) Immunoblotting of UNC-45A, NM-IIA and NM-IIC expression in isolated human colonic epithelium of patients with Crohn’s disease, ulcerative colitis and non-IBD controls. Representative immunoblots (B) and densitometric quantification (C) of UNC-45A expression are shown. Means ± SE (n=6); **P< 0.005. Scale bar, 20 μm.

**Supplemental Figure 3. UNC-45A interacts with NM-IIA in polarized intestinal epithelial cell monolayers.**

(A) Immunoprecipitation analysis of UNC-45A-NM-II interactions in SK-CO15 cells. UNC-45A was pulled-down with anti-UNC-45A antibody. Immunoprecipitation with IgG is added as a control. Immunoprecipitates were probed for UNC-45A and NM-IIA.

(B,C) Proximity ligation assay using a pair of UNC-45A and NM-IIA antibodies in control and UNC-45A-depleted IEC. Representative fluorescence microscopy images (B) and quantification of the number of dots indicating interactions are shown. For the negative control, the NM-IIA antibody was omitted. Means ± SE (n=4); ***P< 0.0005. Scale bar, 20 μm. A-C data are representative of two independent experiments.

**Supplemental Figure 4. Loss of UNC-45A attenuates proliferation of SK-CO15, but not HT-29 cells without causing cell apoptosis.**

(A,B) MTT assay of control and UNC-45A-depleted HT-29cf8 (A) and SK-CO15 (B) cells at different times post-plating. Means ± SE (n=3); *P< 0.05, **P< 0.01.

(C,D) Immunoblotting analysis of cleaved caspase 3 and cleaved PARP in total cell lysates of control and UNC-45A-depleted HT-29cf8 (C) and SK-CO15 (D) cells. Staurosporine treatment was used as a positive control for apoptosis induction. A-D data are representative of three independent experiments.

**Supplemental Figure 5. UNC-45A regulates assembly of focal adhesions and organization of the actin cytoskeleton in migrating intestinal epithelial cells.**

(A,B) Immunofluorescence labeling of a focal adhesion marker, phosphorylated (p) paxillin, at the leading edge of wounded control and UNC-45A-depleted SK-CO15 cell monolayers. Representative confocal microscopy images (A) and quantification of the focal adhesion number (B) are shown. Each dot represents the average number of focal adhesion counted in three different images. Means ± SE (n=4); **P< 0.005. Scale bar 20 μm.

(C) Confocal microscopy images of F-actin labeling at the leading edge of migrating control and UNC-45A-depleted SK-CO15 cell monolayers. Scale bar, 20 μm. A-C data are representative of three independent experiments.

**Supplemental Figure 6. Loss of UNC-45A does not affect tensile forces at E-cadherin complexes localized at intercellular contacts.**

(A,B) FRET analysis of control and UNC-45A knockout HT-29cf8 cells expressing an E-cadherin FRET-based biosensor. Representative images (A) and the scatter dot plot (B) shows FRET index measurements across monolayers. The horizontal line (black) in the scatter dot plot denotes the median of the FRET index estimates and the whiskers denote the largest observation within 1.5 x interquartile range (IQR) (representative data shown is from one experiment out of a total of four independent experiments; N.S, not significant, p>0.05, One-Way ANOVA and post-hoc Tukey test. Scale Bar, 20 μm.

**Supplemental Figure 7. UNC-45A depletion does not alter organization of apical microtubules.**

Confocal microscopy images of control and UNC-45A knockout HT-29cf8 (A) and SK-CO15 (B) cells immunolabeled for alpha-tubulin. Arrows point on normal organization of the apical microtubule network in control and knockout cells. Scale bar, 20 μm. Data are representative of three independent experiments, with two different coverslips per condition.

**Supplemental Figure 8. Expression of NM-II interacting N-terminal deletion UNC-45A mutant selectively attenuates intestinal epithelial cell migration.**

Wound healing of control SK-CO15 cell monolayers and cells expressing either GFP-tagged full-length UNC-45A or two different deletion mutants, ΔTPR-GFP (N-terminal truncated), or ΔUCS-GFP (C-terminally truncated). Representative wound images (A) and quantification of wound healing (B) are shown.

Means ± SE (n=6); **P< 0.005. Scale bars, 200 μm. Data are representative of three independent experiments.

## Notes

### Competing Interest Statement

The authors have declared no competing interest.

