## Supplementary figures and images for "A myosin chaperone, UNC-45A, is a novel regulator of intestinal epithelial barrier integrity and repair"

### Supplemental Figure 1

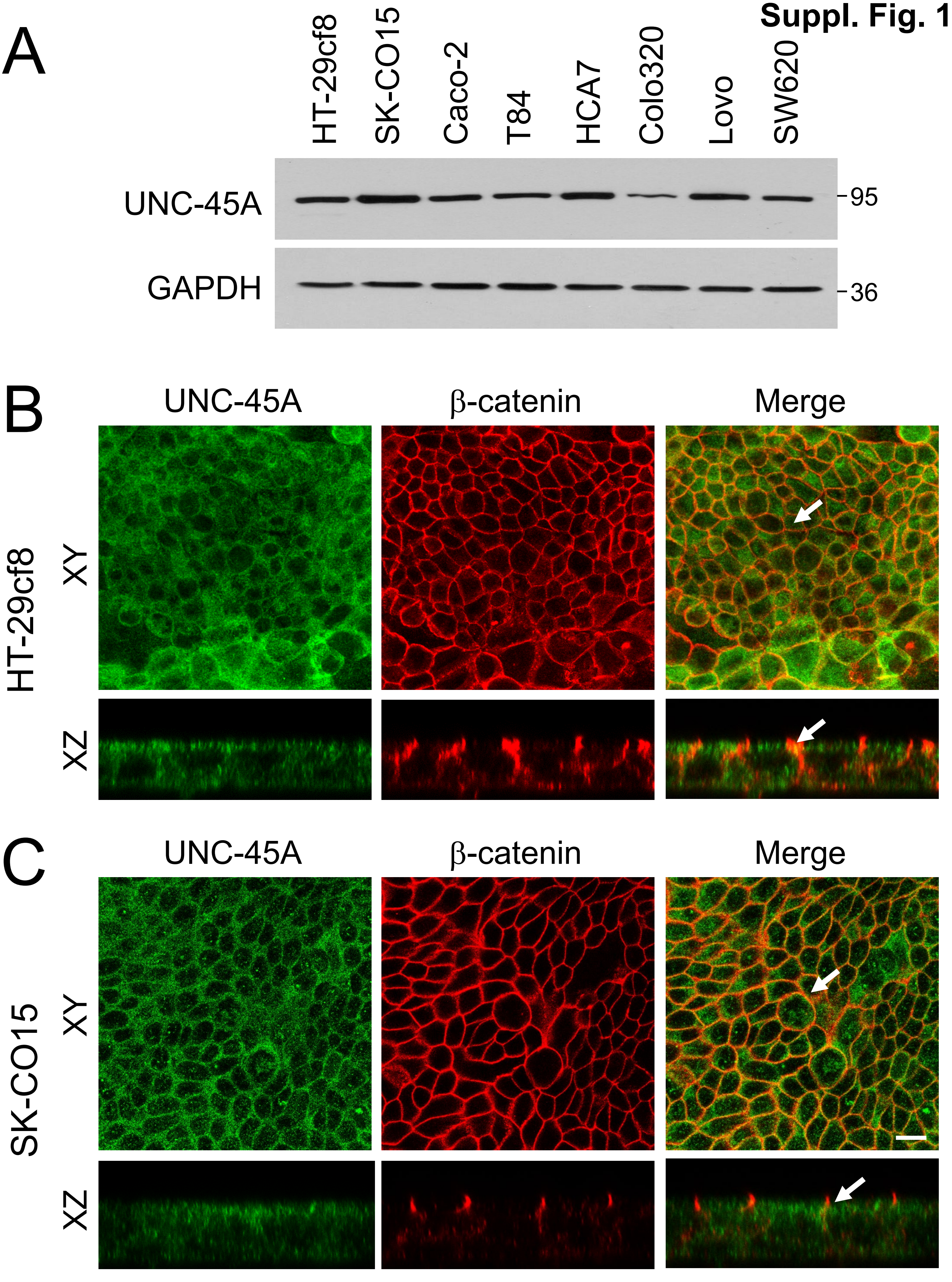

### Supplemental Figure 2

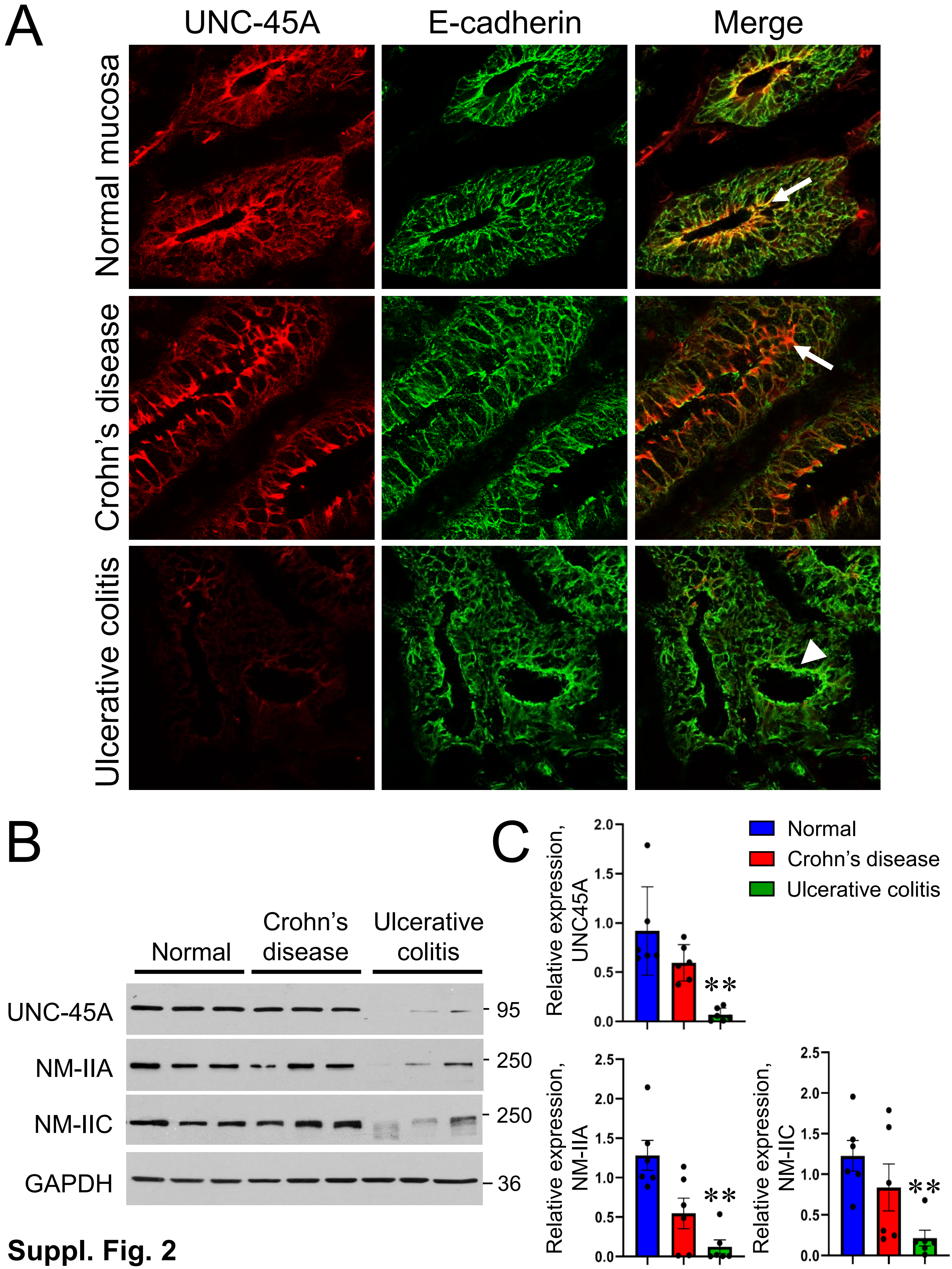

### Supplemental Figure 3

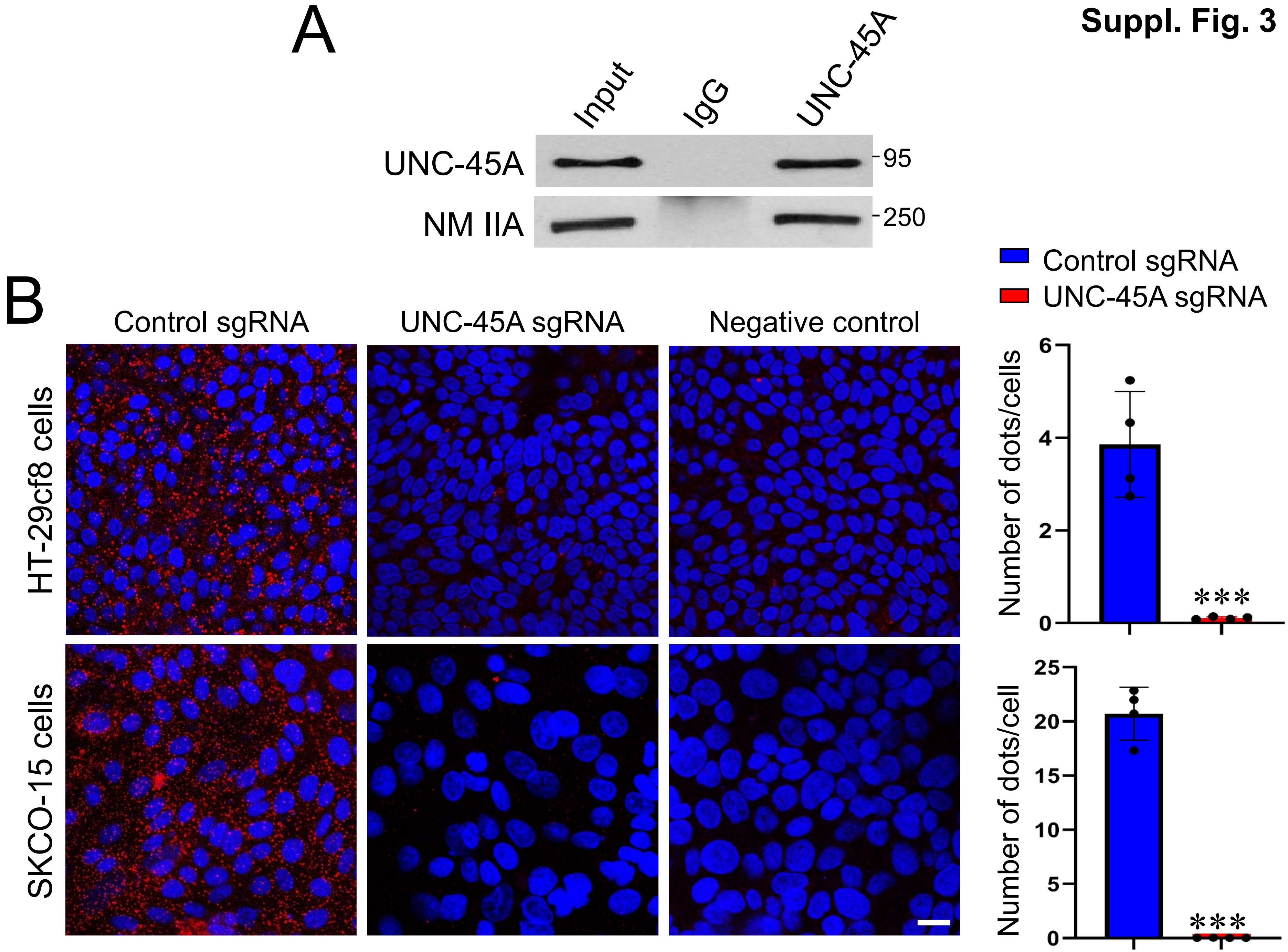

### Supplemental Figure 4

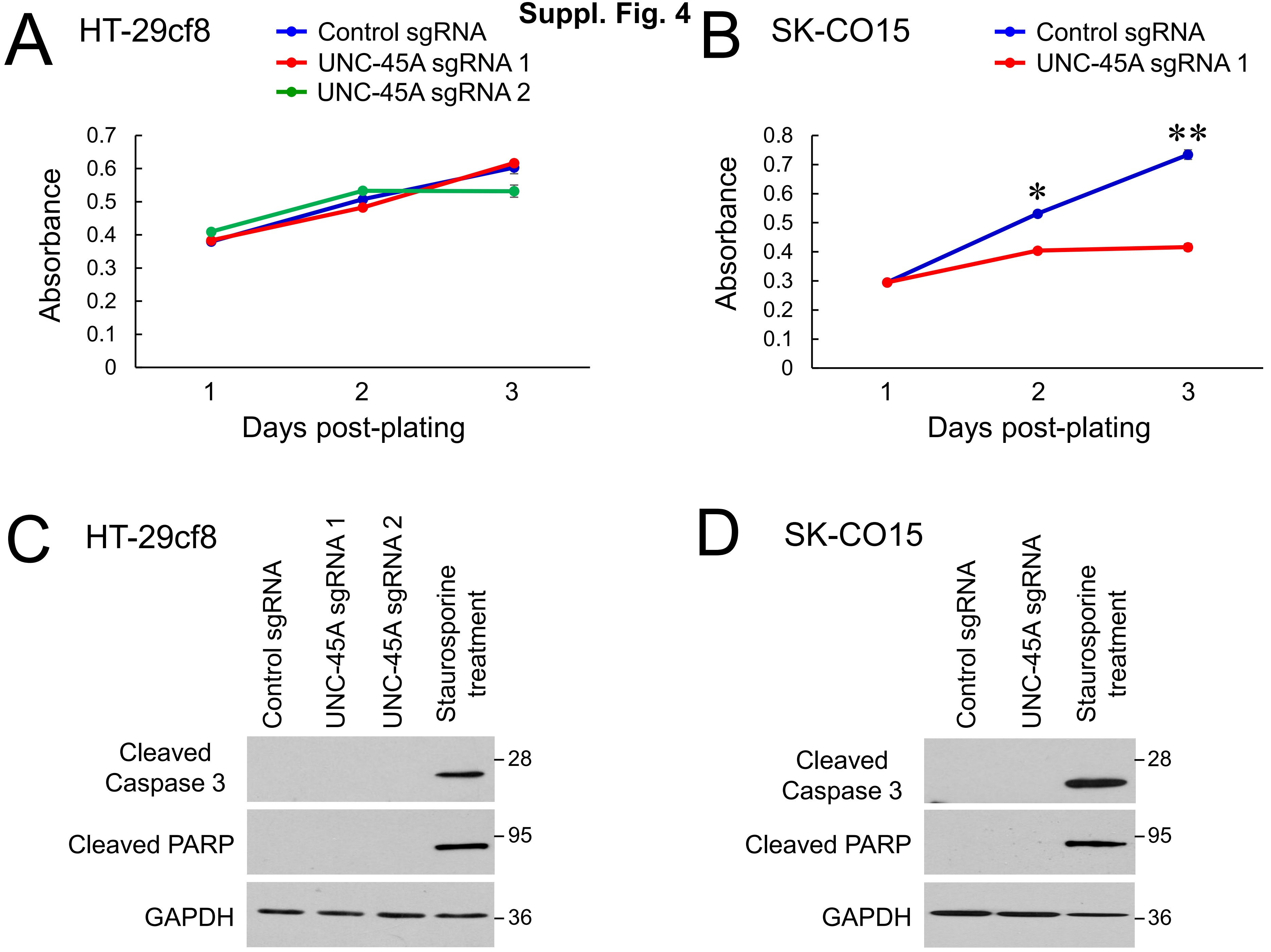

### Supplemental Figure 5

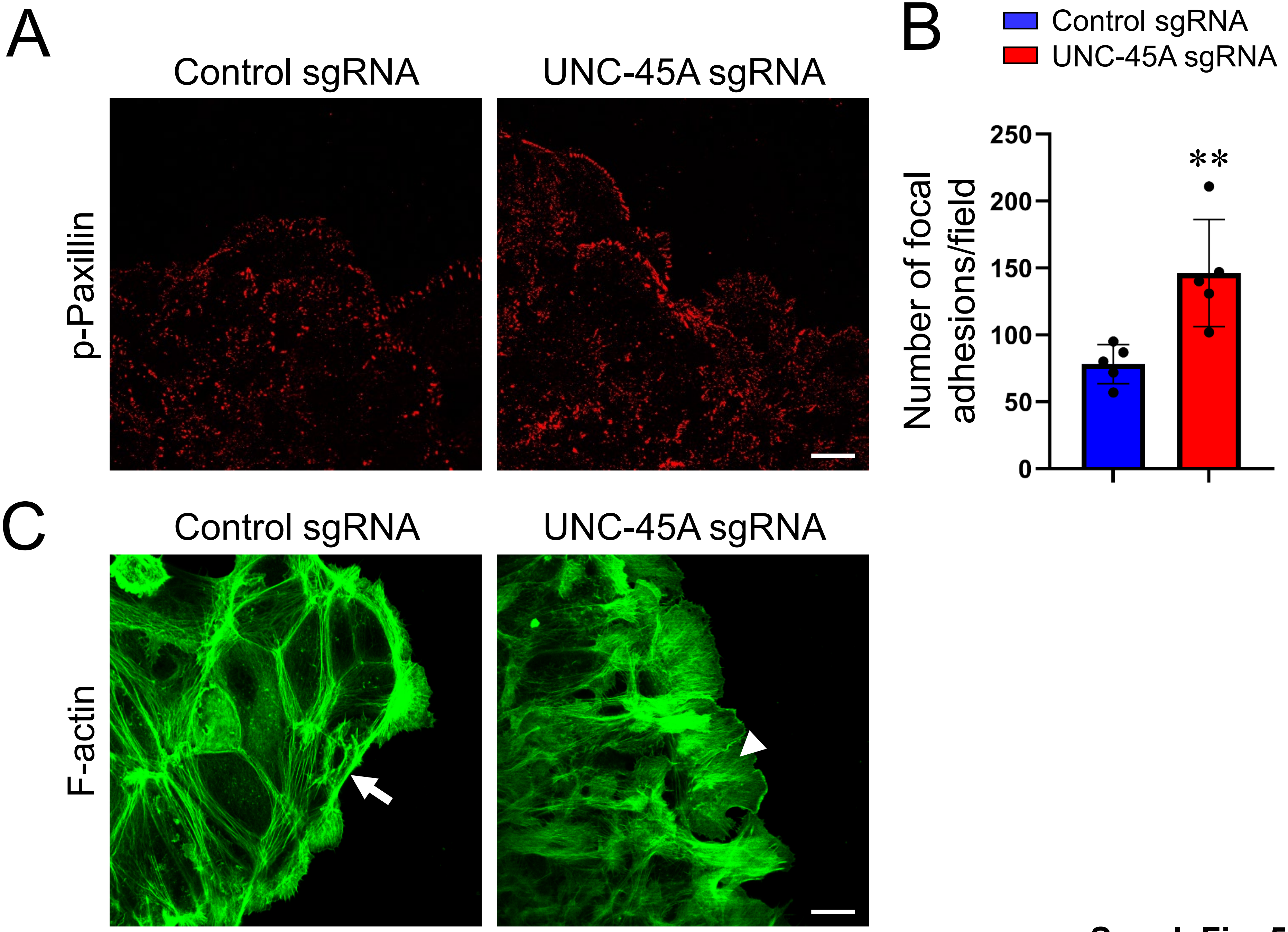

### Supplemental Figure 6

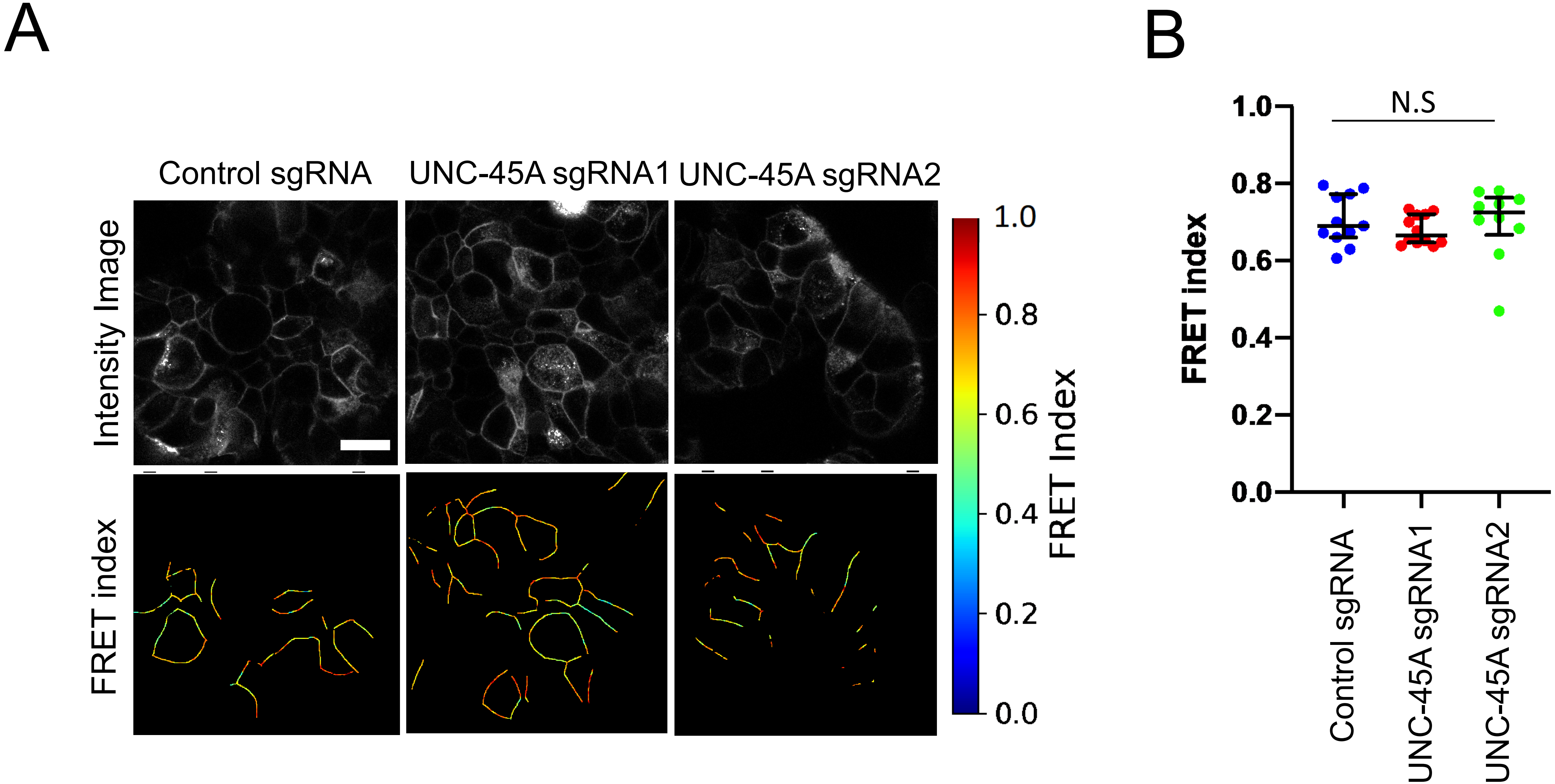

### Supplemental Figure 7

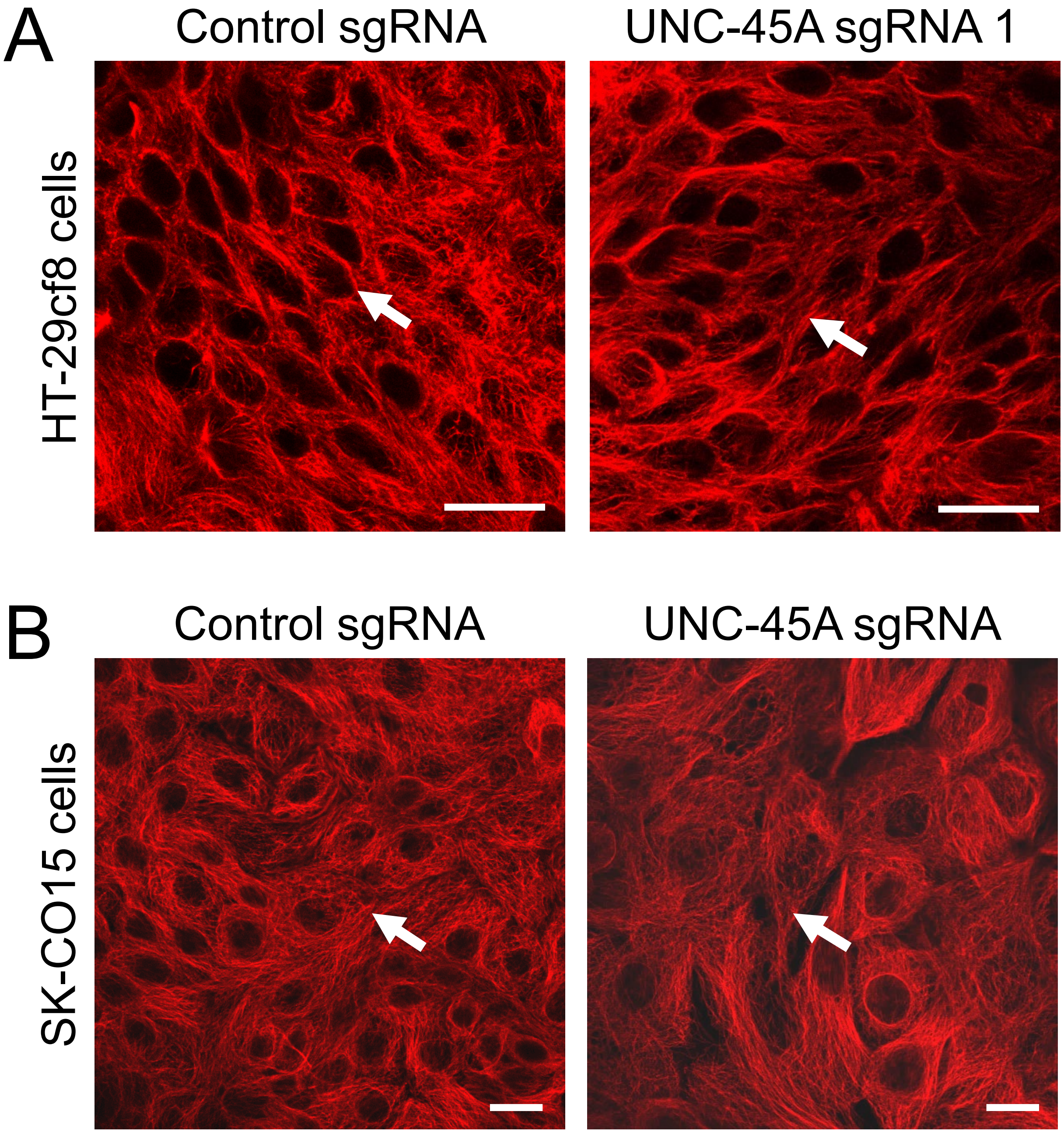

### Supplemental Figure 8

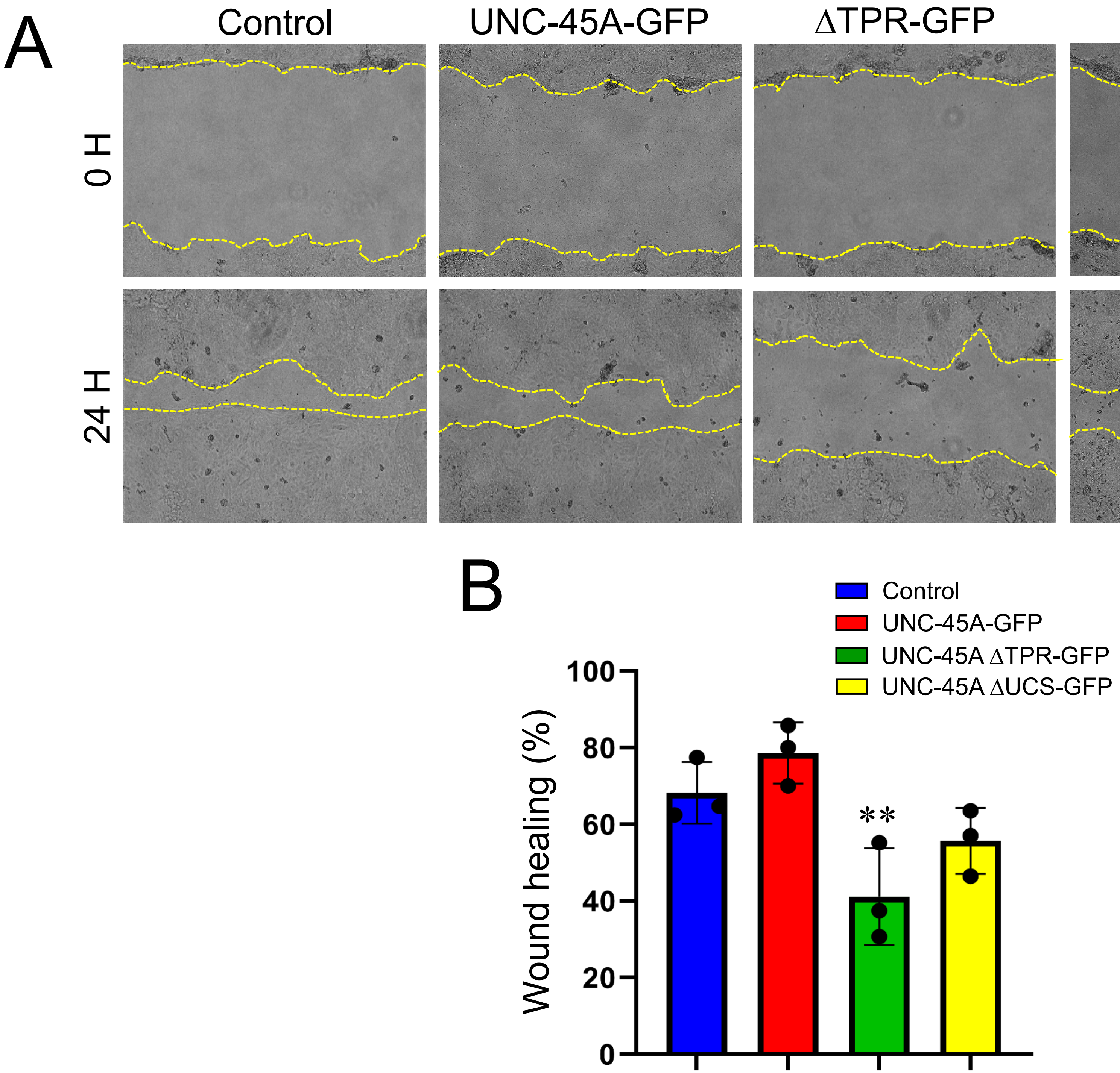
